# Induction of TFEB promotes Kupffer cell survival and reduces lipid accumulation and inflammation in MASLD

**DOI:** 10.1101/2025.04.19.649650

**Authors:** Mandy M. Chan, Sabine Daemen, Wandy Beatty, Kathleen Byrnes, Kevin Cho, Daniel Ferguson, Arick C. Park, Christina F. Fu, Zhen Guo, Natalie Feldstein, Christopher Park, Kira L. Florczak, Li He, Bin Q. Yang, Ali Javaheri, Gary Patti, Brian Finck, Babak Razani, Joel D. Schilling

**Affiliations:** Department of Medicine, Washington University in St. Louis, St. Louis, MO, USA; Department of Medicine, Cardiovascular Research Institute Maastricht, Maastricht University Medical Center, The Netherlands; Department of Molecular Microbiology, Washington University in St. Louis, St. Louis, MO, USA; Department of Pathology and Immunology, Washington University in St. Louis, St. Louis, MO, USA; Department of Chemistry, Siteman Cancer Center, Center for Metabolomics and Isotope Tracing, Washington University in St. Louis, St. Louis, MO 63110, USA; Department of Medicine, Center for Human Nutrition, Washington University in St. Louis, St. Louis, MO 63110, USA; Divison of Cardiology, Massachusetts General Hospital, Harvard Medical School, Boston, MA, USA; Vascular Medicine Institute, Department of Medicine, University of Pittsburgh School of Medicine and UPMC, Pittsburgh, PA, USA

**Keywords:** macrophages, oxidative stress, ferroptosis, lipid peroxidation, *de novo* lipogenesis

## Abstract

Kupffer cells (KCs) are the tissue-resident macrophage of the liver where they serve a critical role in maintaining liver tissue homeostasis and as a filter for circulation. The composition of liver macrophages changes during metabolic dysfunction-associated liver disease (MASLD), with the loss of resident KCs being a hallmark of disease progression. The mechanism(s) and consequences of KC death in metabolic liver disease have yet to be defined. Transcription factor EB (TFEB) is a master regulator of lysosome function and lipid metabolism which has been shown to protect macrophages from lipid stress in atherosclerosis. We hypothesized that TFEB would improve KC fitness in MASLD. To investigate this possibility, we created a transgenic mice in which TFEB was induced specifically in KCs. We found that TFEB induction protected KCs from cell death in two mouse models of MASLD. KC preservation through TFEB induction reduced liver steatosis via a mechanism that was dependent on macrophage lysosomal lipolysis and mitochondrial fatty acid oxidation. The protection from cell death in TFEB KCs was the result of reduced oxidative stress and ferroptosis through a mechanism that involved enhanced NADPH levels. Together, we provide evidence that TFEB promotes KC fitness during MASLD and orchestrates beneficial effects on liver pathology, thus providing potential targets to develop cell-specific therapeutics.

## INTRODUCTION

Metabolic dysfunction-associated steatotic liver disease (MASLD), formerly known as non-alcoholic fatty liver disease (NAFLD), is characterized by excessive fat accumulation in the liver. As MASLD progresses, it is associated with hepatic injury, inflammation, and the formation of fibrotic lesions at which point it evolves into the more severe form known as metabolic dysfunction-associated steatohepatitis (MASH) (*1, 2*). MASLD is the most common form of chronic liver disease, affecting one-third of the population in the Western hemisphere, and MASH-associated cirrhosis is the leading cause of liver transplantation(*3, 4*). Metabolic liver disease is intimately linked with obesity and type 2 diabetes (T2D), and nearly 75% of people with obesity and T2D have MASLD(*5*). Although the FDA recently approved Rezdiffra for the treatment of MASH, only ∼25% of patients treated with Rezdiffra showed an improvement in liver fibrosis(*6, 7*). Therefore, it remains critical to elucidate the cellular and molecular determinants that promote MASLD progression.

Kupffer cells (KCs) are embryonically derived, liver-resident macrophages that account for > 90% of the macrophages in a non-diseased liver(*8, 9*). KCs are located in the lumen of liver sinusoids with a greater density in the periportal area(*10*). These resident macrophages serve important physiological functions such as filtering the systemic circulation, modulating immune activation, and ingesting dead cells and excessive iron(*11–15*). In a normal liver, KCs maintain their cell number by self-renewal; however, loss of KCs by genetic depletion(*16–18*), clodronate-containing liposomes(*10, 19*), or irradiation(*20*) leads to rapid infiltration of monocytes into the liver where they can differentiate into monocyte-derived macrophages (MdMs), some of which fill the niche of the depleted KCs(*16, 17, 19, 20*). Canonical macrophage markers such as F4/80 and CD11b fail to highlight the diversity of macrophages that appear in MASLD; however, the use of single-cell RNA sequencing (scRNAseq), multimarker flow cytometry, and spatial imaging have revealed significant heterogeneity and dynamic changes in the composition of liver macrophages in human and mouse MASLD (*21–25*). Based on these data it is now recognized that KCs express lineage-specific markers such as TIM4, MARCO, CD163, CLEC4F, and VSIG4 that distinguish them from other disease-associated MdMs. Furthermore, we and others have demonstrated that the number of *bona fide* KCs decreases over time in various mouse models of MASLD(*26–29*). While it has been established that hepatocytes accumulate toxic lipid species and succumb to cell death during MASLD (*30–32*), the mechanisms accounting for the loss of resident KCs have been less clear. Recent reports have linked lysosomal stress and iron accumulation to KC death during MASLD (*33, 34*). Given the crucial biological role of KC in metabolizing excessive lipids, clearing dying hepatocytes, and sensing immunological stimuli, it is thus of interest to investigate the potential of reprogramming the native metabolic pathways of KCs and its sequelae.

<u>T</u>ranscription factor <u>EB</u> (TFEB) is a member of the MiT family of transcription factors and serves as a master regulator of lipid metabolism, lysosomal biogenesis, and autophagy(*35–40*). Activation of TFEB has been identified as an attractive therapeutic approach for various diseases involving lysosomal dysfunction, including neurodegenerative diseases(*41*), cerebral ischemia(*42*), and atherosclerosis(*40, 43, 44*). Both genetic overexpression and pharmacological activation of TFEB in macrophages have been shown to 1) rescue lipid-induced lysosome dysfunction and reduce disease severity in atherosclerosis and 2) protect against high-fat diet-induced obesity via growth differentiation factor 15 (GDF15) expression (*43, 44*). In the case of MASLD, TFEB is uniquely positioned to enhance lipid catabolism and lysosome function in a coordinated fashion to promote macrophage fitness. To explore the impact of TFEB activation in KCs during MASLD, we generated mice in which TFEB was overexpressed in a KC-specific manner and assessed the phenotype of these mice on two distinct MASH diets. We discovered that TFEB-overexpression rescues KC death *in vivo*, augments their lipid uptake and metabolism, and reduces liver steatosis after MASLD induction. Mechanistically we demonstrated that TFEB protects macrophages from oxidative stress and cell death through increasing intracellular NADPH levels.

## RESULTS

### Induction of TFEB in Kupffer cells alters macrophage lipid-handling

Increasing TFEB expression or activity has been shown to prevent lipotoxic cell death in macrophages treated with fatty acids (FAs) or cholesterol(*40, 43, 45*). However, these data were largely derived from *in vitro* studies using peritoneal or bone marrow-derived macrophages (BMDMs). To investigate the impact of TFEB on KCs during MASLD, we generated KC-specific TFEB-overexpressing mice (KC^Tfeb^) by crossing the Clec4f^Cre^ line (KC^Cre^)(*17*) with mice containing flox/stop/flox cassette followed by a *Tfeb*-3xFLAG transgene(*39*) (Fig. 1A). To confirm the efficiency of the Clec4f^Cre^ system we demonstrated that *Tfeb* mRNA levels were increased in purified KCs from KC^Tfeb^ mice compared to KC^Cre^ animals. *Tfeb* was not overexpressed in BMDMs from the KC^Tfeb^ mice (Fig. 1B, C). Further qPCR showed that a subset of genes related to lysosome function and lipid metabolism were upregulated in TFEB-overexpressing KCs, whereas the mRNA levels of autophagic genes were not impacted (Fig. 1D). To further confirm the Cre specificity in the setting of MASLD, we crossed these animals to mice containing a *Rosa26*^flox/stop/flox^ TdTomato (TdT) reporter (KC^Cre-TdT^ and KC^Tfeb-TdT^) and fed the mice a high fat, high sucrose, and high cholesterol (HFHS) diet for 16 weeks. We found that over 80% of TIM4^+^VSIG4^+^ KCs were TdT^+^ (fig. S1A-B). MdMs in MASLD consist of monocyte-derived KCs (MoKCs) that have acquired some KC markers (i.e. VSIG4, CLEC4F) and lipid-associated macrophages (LAMs)/pre-MoKCs which are F4/80^hi^ but lack both TIM4 and VSIG4 expression and frequently express *Trem2* (fig. S1A)(*21, 26, 27*). In our reporter mouse, we found that 60-80% of KC precursor MoKCs (TIM4^-^VSIG4^+^) were TdT^+^, but LAMs and monocytes were mostly reporter-negative (fig. S1B).

**Figure 1.**
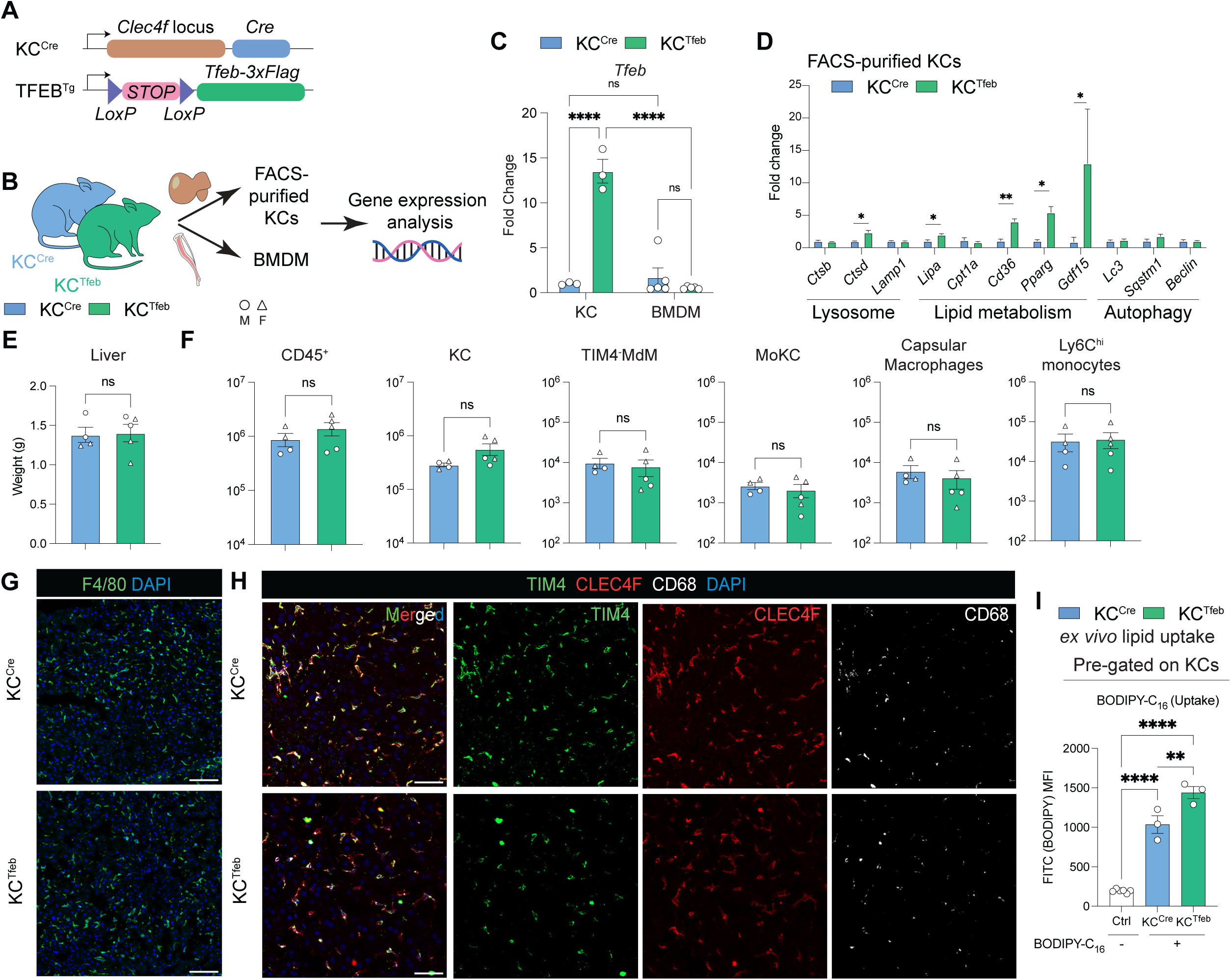
Overexpression of TFEB in KCs enhances lipid uptake and metabolic gene expression. (**A**) Constructs to generate KC-specific TFEB-overexpressing mice, or KC^Tfeb^. (**B**) Schematics for isolating KCs and BMDMs from KC^Cre^ and KC^Tfeb^ animals for gene expression analyses. (**C**) Gene expression of *Tfeb* in KCs and BMDMs isolated from KC^Cre^ and KC^Tfeb^ mice (n = 3/group; technical duplicate for BMDMs). (**D**) Gene expression of selected targets of TFEB in WT- or TFEB-KCs (n = 3/group). (**E-H**) Characteristics of 8- to 10-week-old KC^Cre^ and KC^Tfeb^ mice. (E) Liver weights and (F) flow cytometric quantification of different macrophage and monocyte populations (n = 4-5/group). Circles represent males; triangles represent females. (G) Representative immunofluorescence images of livers showing overall macrophage distribution. Green: F4/80; Blue: DAPI. Scalebar = 100µm. (H) Representative immunofluorescence images of livers showing KC-specific markers with pan-macrophage markers CD68. Green: TIM4; Red: CLEC4F; White: CD68; Blue: DAPI. Scalebar = 50µm. (**I**) Mean fluorescent intensity (MFI) of BODIPY-C_16_ signal in WT or TFEB-KCs quantified by flow cytometry (n = 3/group). Ctrl = no BODIPY-C_16_ controls. Data represent individual biological replicates and are presented as means ±SEM. P-values were calculated using (C) two-way ANOVA followed by multiple t-tests were performed, (D-E) unpaired two-tailed Student’s t-tests were performed, and (I) one-way ANOVA followed by multiple t-tests were performed. NS = not significant, *p < 0.05, **p< 0.01, ***p<0.001, ****p <0.0001.

At baseline, TFEB induction in KCs did not change liver weight, the number of KCs, MdMs, or Ly6C^hi^ monocytes when compared to control mice (Fig. 1E, F, fig. S1A). Moreover, the localization, morphology, and cell surface marker expression of KCs were similar between the genotypes as determined by immunofluorescence and flow cytometry (Fig. 1G, H). Functionally, overexpression of TFEB in KCs increased the expression of *Cd36* and enhanced FA uptake *ex vivo* (Fig. 1D, I). Nevertheless, micropinocytosis of dextran and lysosomal number/activity appear similar between WT and TFEB-KCs at baseline (fig. S1C-D), which may reflect the high baseline lysosomal activity of KCs. Thus, we conclude that TFEB-overexpression in KCs promotes the uptake of FA without affecting baseline macrophage number and composition in the liver.

### KC-specific TFEB induction preserves KC number and reduces MdM infiltration

To assess the impact of augmenting TFEB in KCs on MASLD pathogenesis, we placed male and female KC^Cre^ and KC^Tfeb^ mice on HFHS diet for 16 weeks (Fig. 2A). Male mice of both genotypes gained similar degrees of weight with comparable increases in liver, gonadal adipose tissue, and spleen size after 16 weeks of HFHS diet (Fig. 2B). As expected, female mice did not gain as much weight, but again there was no difference between the genotypes (Fig. S2A). The degree of glucose intolerance was also similar between the genotypes after HFHS diet (fig. S2B).

**Figure 2.**
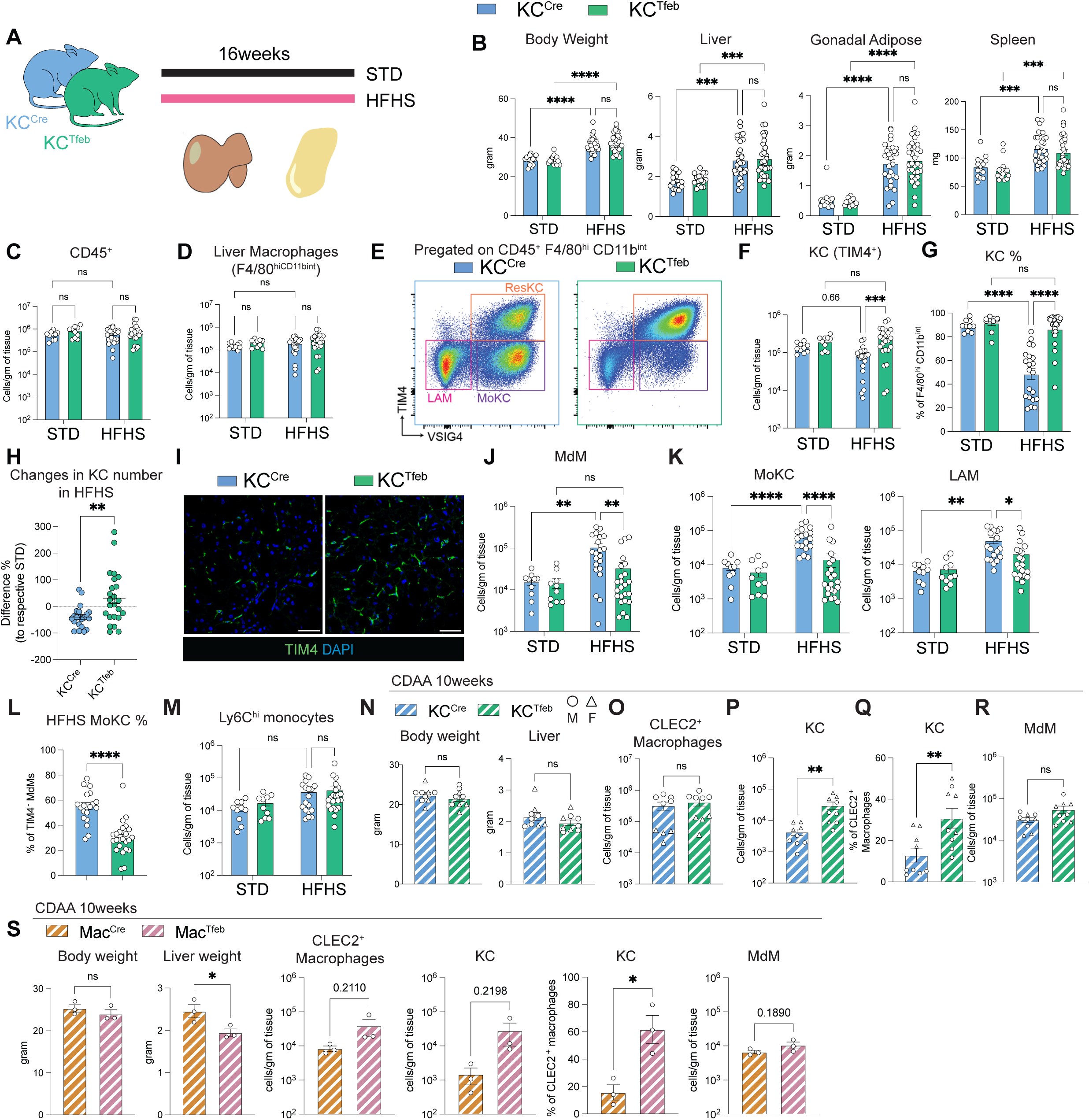
TFEB induction maintains KC numbers and reduces MdM accumulation in MASLD. (**A-M**) STD vs. HFHS diet experiments with 8-10-weeks-old male KC^Cre^ and KC^Tfeb^ littermate mice (n = 10-24/group). (A) Schematic of experiment. (B) Final body and organ weights of STD-fed or HFHS diet-fed mice. (C) Flow cytometric quantification of live, single CD45^+^ cells, (D) total liver macrophages (CD45^+^F4/80^hi^CD11b^int^). (E) Representative flow plots of liver macrophages in HFHS livers. (F) Flow cytometric quantification of KCs (TIM4^+^). (G) Percentage of KCs within total liver macrophages. (H) Changes in KC numbers comparing HFHS-fed KC^Cre^ and KC^Tfeb^ littermates with respective STD-fed mice. (I) Representative immunofluorescence images of livers of HFHS diet-fed mice stain with TIM4 to identify KCs. Green: TIM4; blue: DAPI. Scalebar = 50µm. (J) Flow cytometric quantification of MdMs (TIM4^-^) and (K) MdM subsets MoKCs (TIM4^-^VSIG4^+^) and LAMs (TIM4^-^VSIG4^-^). (L) Percentages of MoKCs among total TIM4^-^ MdMs. (M) Flow cytometric quantification of Ly6C^hi^ monocytes (F4/80^lo^CD11b^+^MHCII^-^Ly6C^+^). (**N-R**) KC^Cre^ and KC^Tfeb^ littermates were fed 10 weeks of fibrogenic CDAA diet (n = 9/group). Circles represent males and triangles represent females. (N) Final body and liver weight. (O) Flow cytometric quantification of CLEC2^+^ liver macrophages (CLEC2^+^F4/80^hi^CD11b^int^) and (P) KCs (TIM4^+^). (Q) Percentage of KCs in CLEC2^+^ macrophages. (R) Quantification of MdMs (CLEC2^+^TIM4^-^) (**S**) Body weights, liver weights, quantification of KC number and percentages, and quantification of MdMs in male Mac^Cre^ and Mac^Tfeb^ littermates fed 10 weeks of CDAA diet (n = 3/group). Data represent individual biological replicates and are presented as means ±SEM. P-values were calculated using (A-D, F-G, J-M) two-way ANOVA followed by multiple t-tests and (H, N-S) two-tailed unpaired t-tests. NS = not significant, *p < 0.05, **p< 0.01, ***p<0.001, ****p<0.0001.

To determine the effect of TFEB on KC survival, we conducted flow cytometry on the livers of male KC^Cre^ and KC^Tfeb^ mice after diet. Neither diet nor genotype changed the number of CD45^+^ leukocytes or overall liver macrophages (F4/80^hi^CD11b^int^) (Fig. 2C, D). However, while KC number began to decline in control mice on HFHS diet, TIM4^+^ KC numbers were maintained in KC^Tfeb^ mice (Fig. 2E-H). These findings were also confirmed by immunofluorescence imaging (Fig. 2I). Loss of KCs can trigger the recruitment and accumulation of MdMs in the liver (*16, 17, 19, 20, 26–28*). KC^Tfeb^ mice had fewer MdMs in the liver following HFHS diet feeding (Fig. 2E, J). Although both MoKCs and LAMs were decreased in the KC^Tfeb^ livers, MoKCs were more dramatically reduced, consistent with the greater preservation of resident KCs in these mice (Fig. 2K, L). The number of Ly6C^hi^ monocytes and lymphocyte populations also did not differ significantly between the genotypes (Fig. 2M, fig. S2C).

We recently reported that steatosis can drive infiltration of MdMs to the liver independent of KC loss(*46*). To determine whether TFEB in KCs impacts the early stage of MdM recruitment we placed KC^Cre^ and KC^Tfeb^ mice on a short-term (8-week) HFHS diet to induce mild liver steatosis (fig. S2D). At this time point, mature KC numbers were preserved in both WT and transgenic mice; however, we still found a reduction in mature MdMs number in the KC^Tfeb^ livers (fig. S2E-I). Similar findings were seen in female mice fed 16 weeks of HFHS diet even though they did not gain as much weight as their male counterparts (fig. S2J-O).

To further validate that TFEB protects KCs in MASH we fed KC^Cre^ and KC^Tfeb^ mice a highly inflammatory, non-obesogenic choline-deficient amino acid-defined (CDAA) diet for 10 weeks to induce significant steatosis and fibrosis. Body weight and liver size were again similar between the genotypes (Fig. 2N). As this diet induces robust infiltration of monocytes and immature MdMs, we utilized CLEC2 positivity to identify mature liver macrophages(*28*) (fig. S2P). CLEC2^+^ liver macrophage numbers were similar for both genotypes on diet, and similar to the HFHS diet TFEB-KCs persisted better than WT control KCs (Fig. 2O-Q). The number of MdMs markedly increased with CDAA diet and was similar between the genotypes (Fig. 2R). To ensure the robustness of this observation, we repeated the diet experiment with mice with LysM-Cre (Mac^Cre^) crossed to the TFEB transgene to drive TFEB overexpression in all macrophages (Mac^Tfeb^). Similar to the findings with the Clec4f-Cre (KC^Cre^) system, KC number was higher in Mac^Tfeb^ mice after CDAA diet feeding (Fig. 2S). Thus, we conclude that TFEB induction maintains resident KC numbers and reduces MdM recruitment in MASH.

### TFEB induction in KCs alters the hepatic lipid landscape in MASLD

We next investigated the impact of TFEB induction in KCs on liver pathology associated with MASLD. To evaluate hepatic steatosis, we first performed histologic assessment of liver tissue. H&E sections were scored by a blinded pathologist and revealed that KC^Tfeb^ mice had a reduction in macrovesicular, but not microvesicular, steatosis in the liver (Fig. 3A-C). In addition, liver triglyceride (TAG) and plasma alanine transaminase (ALT) levels were modestly reduced in the KC^Tfeb^ mice while liver cholesterol and plasma triglyceride concentration remained unchanged (Fig. 3D-E, fig, S3A). The expression of *de novo* lipogenesis or fatty acid oxidation (FAO)-related genes in whole liver was significantly increased by HFHS diet but was not different between genotypes (Fig. 3F). To further quantify the lipid changes we conducted targeted lipidomics for several TAG species (fig. S3B). HFHS feeding was associated with a significant increase in most TAG species when compared to STD-fed livers (Fig. 3G). Among the TAG species measured, those with 16 carbons, particularly unsaturated fatty acids (X:1 or X:2) were significantly reduced in the KC^Tfeb^ livers despite similar liver weights (Fig. 3G, H, fig. S3B). Livers from KC^Tfeb^ mice had increased levels of low abundance TAG containing unsaturated 18 carbon chains at baseline (fig. S3C). More severe MASH also leads to liver fibrosis; however, the extent of fibrosis in HFHS diet was minimal (fig. S3D). Therefore, we performed picrosirius red (PSR) imaging on the livers of KC^Cre^ and KC^Tfeb^ mice after 10 weeks of the fibrogenic CDAA diet. While 4 out of 5 KC^Tfeb^ livers imaged have lower PSR area, the average extent of fibrosis by PSR quantification was similar between the genotypes (fig. S3E). Taken together, KC preservation and/or induction of TFEB in KCs is sufficient to alter the lipid landscape in the liver without directly changing hepatocyte lipid metabolic programs or stellate cell fibrotic response in MASLD.

**Figure 3.**
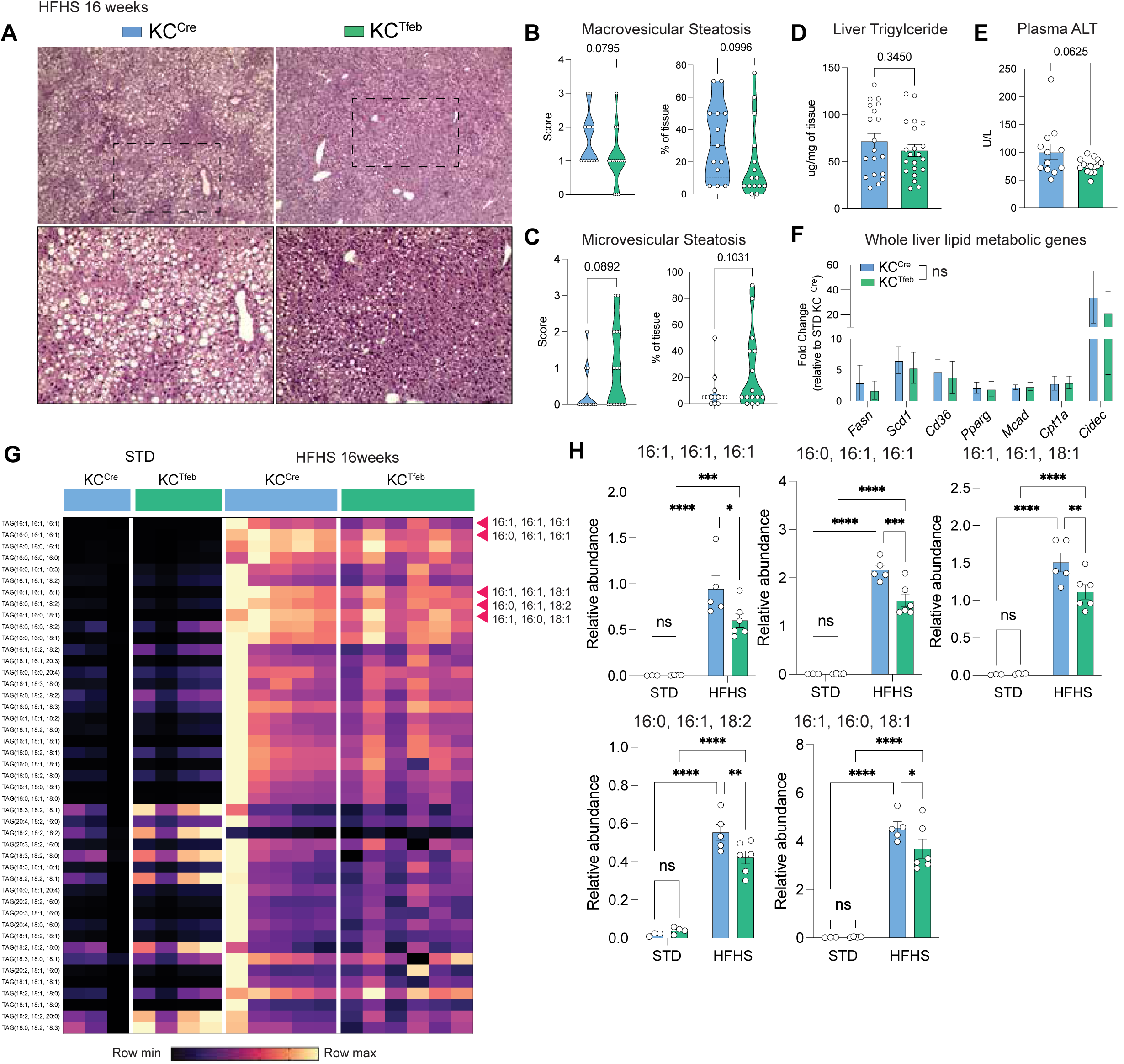
TFEB induction in KCs reduces hepatic steatosis. (**A-D**) Liver steatosis phenotyping for 16-week HFHS diet-fed KC^Cre^ and KC^Tfeb^ mice (n = 13-21/group). (A) Representative H&E images of livers. (B) Macrovesicular and (C) microvesicular steatosis scores and percentages as determined by a blinded pathologist. (D) Liver TAG measurement. (**E**) Plasma ALT measurement from 16-week HFHS diet-fed KC^Cre^ and KC^Tfeb^ mice (n = 12-15/group). (**F**) qPCR gene expression analyses of selected lipid metabolic genes in whole liver tissues of KC^Cre^ and KC^Tfeb^ mice after 16 weeks of STD or HFHS feeding (n = 5-7/group). Shown are fold changes relative to STD-fed KC^Cre^. (**G**) Heatmap of normalized (within individual TAG species) relative abundance of 45 TAG species from hepatic tissue, accompanied by quantification of selected species with statistical significance (n = 3-6/group). Data represent individual biological replicates and are presented as means ±SEM. P-values were calculated using (B-F) unpaired two-tailed Student’s t-tests, and (H) two-way ANOVA followed by multiple t-tests. NS = not significant, *p < 0.05, **p< 0.01, ***p<0.001, ****p<0.0001.

### MASLD induces lysosomal, metabolic, and pro-survival transcriptomic signatures in TFEB-overexpressing KCs

To unravel potential mechanisms by which TFEB expression preserves KC number and alters hepatic lipid accumulation in MASLD, we performed bulk RNA sequencing on flow-sorted TIM4^+^ KCs from KC^Cre^ and KC^Tfeb^ animals fed STD or HFHS diet for 16 weeks (Fig. 4A). At baseline, TFEB induction in KCs led to upregulation of genes related to cholesterol metabolism and lipoic acid metabolism (fig. S4A, B). In response to HFHS diet TFEB-KCs significantly diverge from WT-KCs in their transcriptional response (fig. S4C). After HFHS feeding, the upregulated differentially expressed genes (DEGs) in WT-KCs mapped to PPAR signaling pathway while downregulated DEGs linked to pathways related to IL-17 and MAPK signaling (fig. S4D, E). In TFEB-KCs, HFHS diet led to the upregulation of pathways related to lysosomes, ER protein processing, and PPAR signaling as well as downregulation of amino acid metabolism (fig. S4F, G). To discern the effects of TFEB and the diet on KC phenotypes, we compared the significantly upregulated genes in TFEB-KCs in STD and HFHS conditions and identified 70 diet-independent TFEB-modulated genes in KCs (Fig. 4B). Among the 70 genes, 13 of them were related to cholesterol metabolism and lysosomes (Fig. 4C). Outside of the core TFEB-modulated genes, 794 genes were uniquely expressed by HFHS TFEB-KCs vs. 240 by STD TFEB-KCs compared to their WT controls (Fig. 4B). The upregulated DEGs in TFEB-KCs included genes such as *Ctsb*, *Ctss*, *Mcoln2*, *Lipa*, *Gdf15*, *Scd2*, and *Sqstm1* (Fig. 4D). KEGG pathway analysis of these upregulated genes indicated enrichment in metabolic pathways such as lysosome, oxidative phosphorylation, fatty acid elongation, steroid biosynthesis, and mitophagy (Fig. 4D-G). Thus, the combination of TFEB and HFHS diet amplified the differences between genotypes. KC2s are a subset of KCs that are purported to have CD36-mediated metabolic function(*25*). In our RNAseq data most of the KC2-associated transcripts, besides *Cd36,* were either unchanged or downregulated in HFHS TFEB-KCs (fig. S4F). Regarding cell death/survival, a variety of cathepsins as well as anti-apoptosis genes such as *Bcl2a1a-d*, *Bcl2l1*, and *Mcl1* were upregulated in TFEB-KCs (Fig. 4H). Thus, TFEB induction in KCs enhances pathways related to lipid metabolism and lysosome function during metabolic stress.

**Figure 4.**
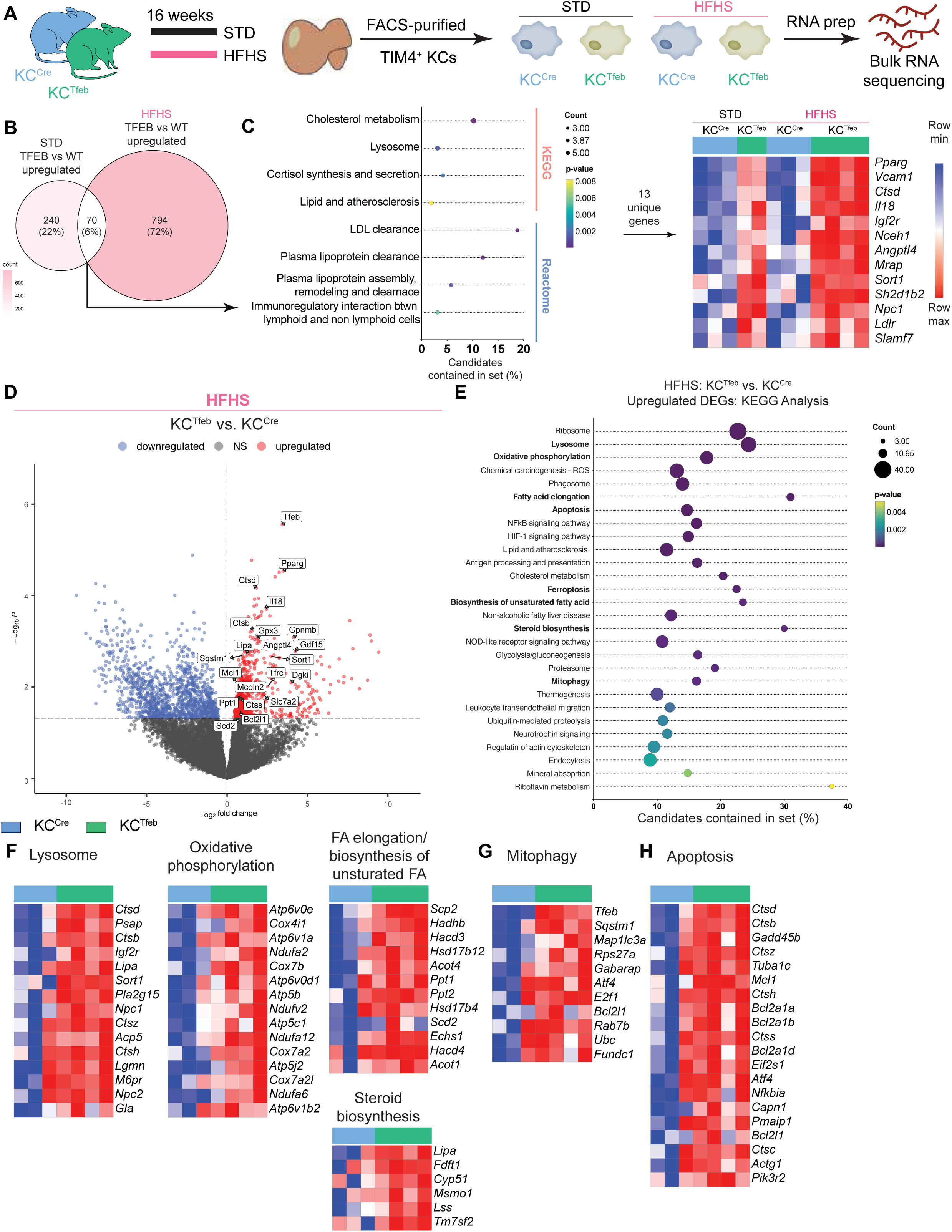
MASLD augments TFEB-mediated lysosomal and lipid metabolic transcriptional programs in KCs. (**A**) Schematic of experiments to isolate KCs from different conditions for bulk RNAseq (n = 2-4/group). (**B-C**) (B) Venn diagram comparing significantly upregulated DEGs and (C) KEGG and Reactome pathway analyses of the 70 commonly upregulated genes and among which 13 unique genes contributed to the pathway analyses in HFHS TFEB- vs. WT-KCs and STD TFEB-vs. WT-KCs. (**D-F**) (D) Volcano plots, (E) KEGG pathways analysis, (F-H) heatmaps of DEGs in TFEB- vs. WT-KCs after HFHS feeding. Log_2_fold change > 0, p-value <0.05.

### TFEB activates lipid metabolic programs in KCs

In WT mice, we found that KCs isolated from 16-week-HFHS-fed mice had increased FA uptake compared to STD-fed mice (fig. S5A); however electron microscopy (EM) of TIM4^+^KCs from a MASLD liver revealed fewer lipid droplets as compared to KCs from STD-fed livers (fig. S5B-E), consistent with our prior findings (*46*). Based on the RNA seq data we hypothesized that altered lipid metabolism in KCs may impact liver steatosis in fatty liver disease. To evaluate this hypothesis, we performed EM on FACS-purified KCs from KC^Cre^ and KC^Tfeb^ mice after HFHS diet. Again, WT-KCs extracted from steatotic livers had few lipid droplets despite the increased lipid content in the liver (Fig. 5A). In contrast, TFEB-KCs had more lipid droplets and lipid droplet area after HFHS diet (Fig. 5A-C). The size distribution of lipid droplets size was also broader in TFEB-KCs (Fig. 5D). Of interest, most droplets in transgenic KCs were in lipid-filled vacuoles (Fig. 5A, E-F). Together these data are consistent with increased capacity for lipid uptake, droplet formation and/or lysosomal lipolysis in TFEB KCs.

**Figure 5.**
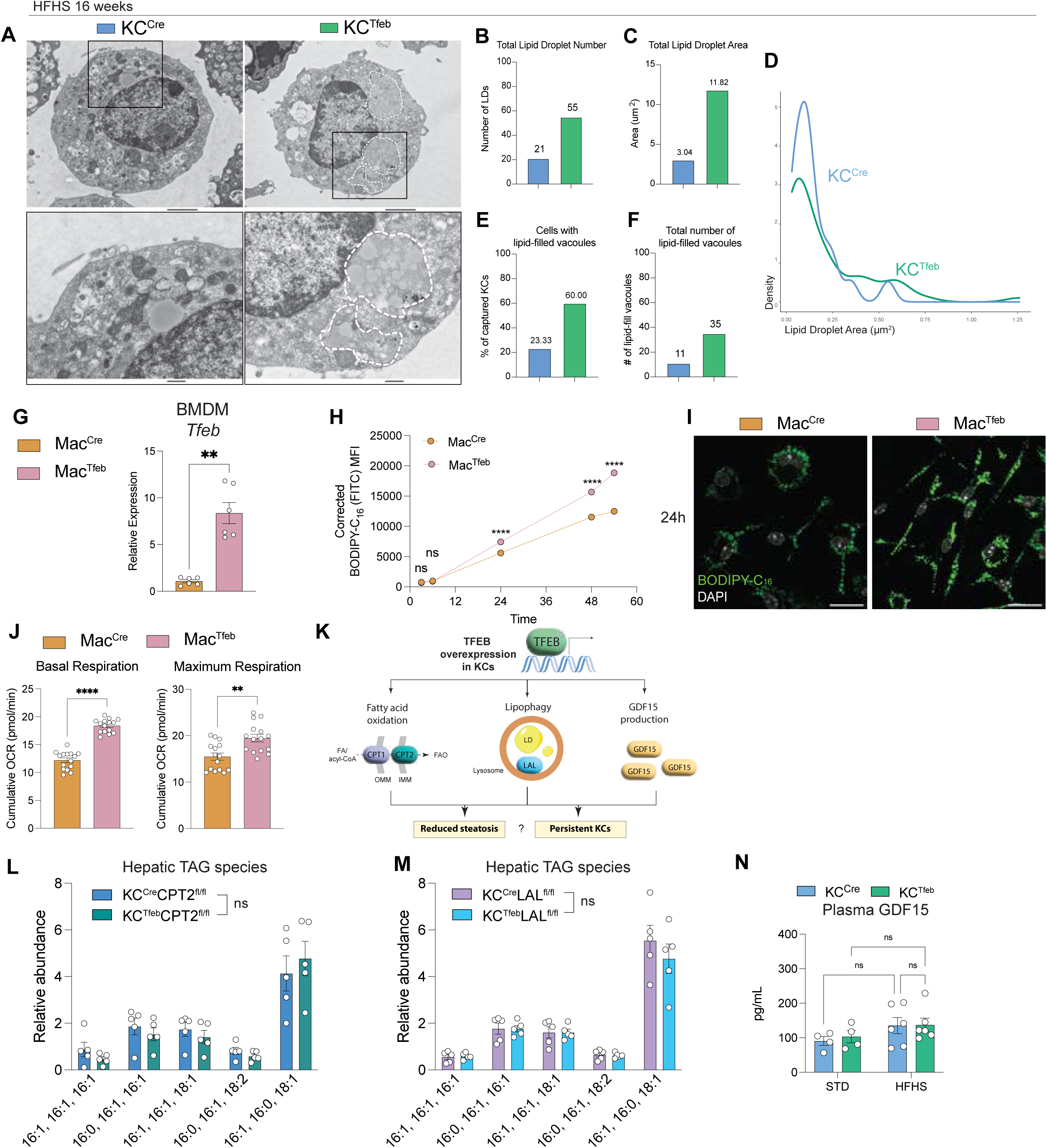
TFEB promotes lipid digestion and oxidative phosphorylation in macrophages. (**A-F**) FACS-purified KCs from HFHS diet-fed KC^Cre^ and KC^Tfeb^ mice were subjected to electron microscopy (n = 2-3/genotype). (A) Representative electron microscopy images of KCs. Top scalebar = 2µm; bottom scalebar = 500nm. (B) Quantification of the total number and (C) total area of LDs across 30 cells per genotype on HFHS diet. (D) Size distribution of LDs per genotype on HFHS diet. (E) Percentage and (F) count of KCs with lipid-filled vacuoles. (**G-J**) *In vitro* experiments using BMDMs derived from Mac^Cre^ and Mac^Tfeb^ mice. (G) qPCR gene expression analysis on *Tfeb* (n = 6/group). (H) MFI corrected for baseline fluorescence and (I) representative immunofluorescence images of BMDMs cultured with 250µM oleic acid conjugated to BSA + 1µM BODIPY-C_16_. Green: BODIPY-C_16_.White: DAPI. Scalebar = 20µm. (J) Cumulative oxygen consumption rate (OCR) for basal and maximum respiration of BMDMs. (K) Graphic representation of potential pathways TFEB induction could alter in lipid handling. (**L-N**) FAO, lysosomal lipolysis, and GDF15-deficient KC^Cre^ or KC^Tfeb^ mice were fed HFHS diet for 16 weeks. (L, M) Hepatic TAG species of KC^Cre^CPT2^fl/fl^, KC^Tfeb^CPT2^fl/fl^, KC^Cre^LAL^fl/fl^, and KC^Tfeb^LAL^fl/fl^ mice (n = 5/group). (N) Plasma GDF15 in KC^Cre^GDF15^fl/fl^, KC^Tfeb^GDF15^fl/fl^ mice (n = 4-6/group), Data represents (G, L-N) individual biological, or (H-J) technical replicates and are presented as (B-F) cumulative data or (G, H, J, L-M) means ±SEM. P-values were calculated using (G) paired and (H, J, L-N) unpaired two-tailed Student’s t-tests. NS = not significant, *p < 0.05, **p< 0.01, ****p<0.0001.

To assess whether TFEB changes the capacity of macrophages to take up and store FAs we utilized BMDMs from Mac^Tfeb^ mice. The degree of *Tfeb* upregulation was similar between the BMDMs prepared from these mice and KCs from the KC^Tfeb^ mice (Fig. 5G, 1C). To address lipid droplet accumulation, we incubated WT or TFEB-BMDMs with BODIPY-labeled C16:0 mixed with unlabeled oleic acid (C18:1) and monitored the accumulation of fluorescent lipid droplets. By 24h TFEB-BMDMs had significantly more lipid droplet accumulation, and this continued out to 54h, a timepoint where the signal in the WT BMDM had begun to plateau (Fig. 5H-I). We also demonstrated that TFEB-BMDMs had greater mitochondrial oxidative capacity using a Seahorse Mitostress test to quantify oxygen consumption (Fig. 5J, fig. S5F). Thus, these observations indicate that TFEB enhances the capacity for lipid storage and fatty acid oxidation in macrophages.

Lipid droplets can be broken down in the lysosome by lysosomal acid lipase (LAL) and these FA can be delivered to the mitochondria via the carnitine palmitoyltransferase (CPT) system for oxidation (*47, 48*) (Fig. 5K). To evaluate the role of KC lysosomal lipolysis and/or mitochondrial oxidation for the attenuated hepatic steatosis phenotype we generated mice in which TFEB overexpression was combined with KC-specific *Cpt2* knockout (KO; KC^Tfeb^CPT2^fl/fl^) to disrupt FAO or KC-specific *Lipa* KO (KC^Tfeb^LAL^fl/fl^) to prevent lysosomal lipolysis. We validated the reduced expression of *Cpt2* and *Lipa* in KCs from these two mouse lines (fig. S5G). After 16 weeks of HFHS diet there was no difference in weight gain or liver TAG levels between any of the mouse lines (fig. S5H, I). The protective effect of KC-TFEB on hepatic steatosis was abolished with CPT2 or LAL deficiency in KCs based on lipidomics and histology (Fig. 5L, M; fig. S5I, K). Thus, the reduction in hepatic steatosis seen in KC^Tfeb^ mice depends on lysosomal degradation and mitochondrial oxidation of lipids in KCs.

Overexpression of TFEB in macrophages has been reported to protect against obesity and insulin resistance via upregulation of GDF15 (*49, 50*) (Fig. 5K). We also found that *Gdf15* mRNA was upregulated in TFEB-KCs (Fig. 1D, 4D). However, in contrast to TFEB overexpression driven by a pan-macrophage LysM^Cre^ system (fig. S5L), GDF15 was not elevated in the plasma of KC^Tfeb^ mice fed HFHS diet (Fig. 5N). To address any potential local effect of KC-produced GDF15, we generated and validated a conditional KO of GDF15 coupled with TFEB-overexpression in KCs (KC^Tfeb^GDF15^fl/fl^) (fig. S5M). After 16 weeks HFHS feeding, both control and KC^Tfeb^GDF15^fl/fl^ mice have similar weight gain and similar levels of hepatic steatosis (fig. S5N-P), indicating that KC-derived GDF15 does not modulate systemic or local metabolism. Together, our results indicate that TFEB induction enhances macrophage lipid handling, allowing for KCs to function as a lipid sink.

### TFEB protects macrophages from cell death largely independent of lysosomal lipid metabolism

The preservation of KCs in KC^Tfeb^ mice in MASLD could reflect increased proliferation, accelerated differentiation of MdMs to KC, or reduced cell death. To assess KC proliferation, we performed Ki67 staining on KCs from KC^Cre^ and KC^Tfeb^ mice after diet and observed no differences between the genotypes (Fig. 6A). We also injected KC^Cre^ and KC^Tfeb^ mice with BrdU after HFHS diet and if anything observed a slight decrease BrdU incorporation in TFEB-KCs (Fig. 6B). Thus, TFEB does not impact KC proliferation during MASH.

**Figure 6.**
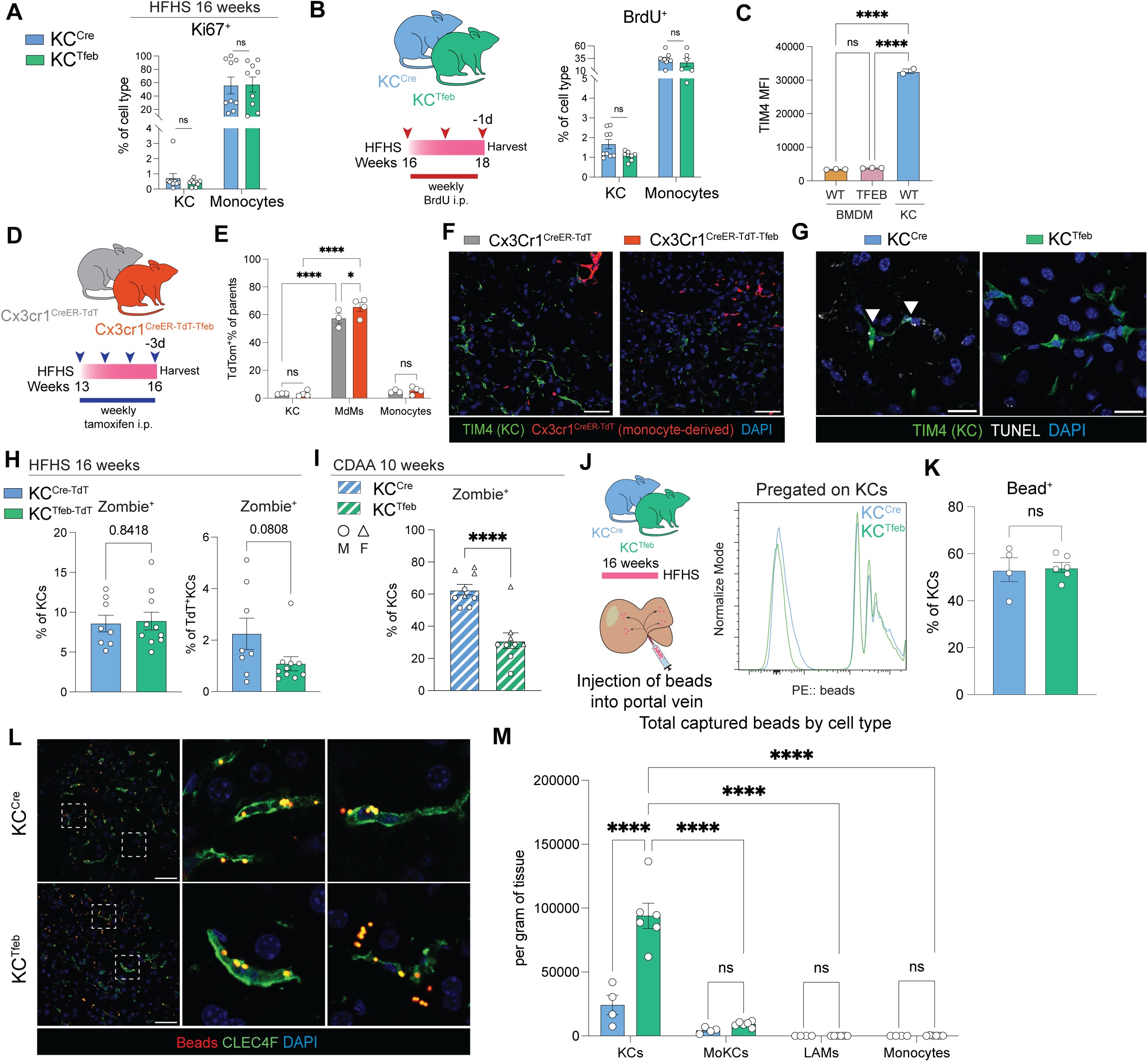
TFEB mitigates macrophage death and improves portal filtration. (**A**) Quantification of Ki67^+^ percentage in KCs after 16 weeks HFHS diet feeding in male KC^Cre^ and KC^Tfeb^ mice (n = 9/genotype). (**B**) Schematic of BrdU incorporation experiment where KC^Cre^ and KC^Tfeb^ littermates fed 16-weeks HFHS diet were intraperitoneally (i.p.) injected with BrdU once a week for 3 weeks, and flow cytometric quantification of BrdU^+^ percentage in cells (n = 8-9/genotype). (**C**) MFI of TIM4-BV421 staining on BMDMs isolated from Mac^Cre^ or Mac^Tfeb^ mice compared to WT-KCs. (**D-F**) Cx3cr1^CreER-TdT^ and Cx3cr1^CreER-TdT-Tfeb^ littermates fed HFHS diet and i.p. injected with tamoxifen once a week for 4 weeks before harvest at 16 weeks (n = 3-4/group) (D) Schematic of injection regimen, (E) flow cytometric quantification of TdT^+^ percentage in cells, and (F) representative immunofluorescence images of livers. Green: TIM4; red: TdT reporter; blue: DAPI. Scalebar = 50µm. (**G**) Representative images of TUNEL staining identified in the livers of 16-week HFHS diet-fed KC^Cre^ and KC^Tfeb^ mice. Green: TIM4; white: TUNEL; blue: DAPI. Scalebar = 20µm. (**H**) Male KC^Cre-TdT^ and KC^Tfeb-TdT^ mice were placed on HFHS diet for 16 weeks and cell death was assessed by Zombie Aqua positivity in total KCs or TdT^+^ KCs (n = 8-10/genotype). (**I**) KC^Cre^ and KC^Tfeb^ mice were placed on CDAA diet for 10 weeks and cell death was assessed by Zombie Aqua positivity in total KCs (n = 9/genotype). Circles represent males; triangles represent females. (**J-M**) KC^Cre^ and KC^Tfeb^ male mice were fed HFHS for 16 weeks and livers were *in situ* injected with fluorescent beads (n = 4-5/group). (J) Schematic of experiment and representative flow histogram of bead signal in KCs. (K) Percentage of KCs with single or multiple bead positive signals. (L) Representative immunofluorescence images bead capturing in KCs. Red: beads; green: CLEC4F; blue: DAPI. Scale bar = 50µm. (M) Total beads captured by liver myeloid cells and monocytes per gram of tissue. Values were determined by multiplying cells of interest per gram of tissue with beads-positive percentage. Data represents individual biological replicates and are presented as means ±SEM. P-values were calculated using (A-B, H-I, K) unpaired two-tailed Student’s t-tests, (C) one-way ANOVA followed by multiple t-tests, and (E, M) two-way ANOVA followed by multiple t-tests. NS = not significant, *p < 0.05, **p< 0.01, ***p< 0.001, ****p<0.0001.

We next evaluated whether TFEB could accelerate the differentiation of MoKCs to TIM4^+^ KCs as MoKCs also express CLEC4F (fig. S1B). First, we found that overexpression of TFEB in BMDMs did not increase resident KC markers such as TIM4 (Fig. 6C). Second, we crossed our TFEB transgenic mice to mice expressing the tamoxifen-inducible Cx3Cr1-CreER system in addition to the TdT reporter (Cx3Cr1^CreER-TdT-Tfeb^). As TIM4^+^ KCs lack expression of *Cx3Cr1*, this system allows for labeling of MdMs with fate mapping to detect TIM4^+^KCs derived from monocytes. Cx3Cr1^CreER-TdT-Tfeb^ and their CreER-only littermates were fed HFHS diet for 12 weeks after which they were injected once a week for 4 weeks with tamoxifen to induce TFEB overexpression and the TdT reporter in MdMs (Fig. 6D). Although we observed robust expression of TdT within MdMs, the reporter was not detected in TIM4 expressing KCs, indicating that TFEB-overexpression does not promote rapid maturation of MdMs into mature KCs (Fig. 6E, F).

To assess the impact of TFEB on macrophage cell death *in vivo* we performed TUNEL staining in conjunction with TIM4 staining and observed fewer dying KCs in KC^Tfeb^ mice after HFHS diet (Fig. 6G). To quantify this effect, we analyzed Zombie staining in a subset of HFHS-fed KC^Cre^ and KC^Tfeb^ mice containing TdT reporter for death signals within KCs. Although total death in KCs was similar between the genotypes, cell death was specifically reduced in TdT^+^KCs, i.e. cells with TFEB induction (Fig. 6H). In mice fed the highly inflammatory CDAA diet, we found a robust reduction of cell death in TFEB-KCs (Fig. 6I). Together, these data argue that the preservation of KCs by TFEB is the consequence of reduced KC death in MASLD.

We also determined whether TFEB loss-of-function also impacts KC survival in MASLD. In contrast to the overexpression system the loss of TFEB in KCs does not affect the survival of these cells, potentially related to compensation by other MiT transcription factor family members such as TFE3 (fig. S6A-D). Although TFEB promotes lysosomal biogenesis, we found that lysosomal activity in WT-KCs exposed to HFHS diet was already increased and that TFEB overexpression did not appear to enhance this further (fig. S6E-G). Therefore, TFEB-mediated KC protection appears to be independent of its effects on lysosomes.

### KC protection by TFEB improves liver filtration in MASLD

KCs are critical in filtering bacteria and particulate from the bloodstream. Therefore, we asked whether the protection of KCs by TFEB might also enhance the capture of particulate antigen. We first established an assay to assess KC filtration function by injecting fluorescent beads into the portal vein of animals that had been fed STD or HFHS diet for 16 weeks (fig. S6H). Using flow cytometry, we quantified bead capture and determined whether KCs ingested one or multiple beads (fig. S6I). After HFHS feeding KCs were less likely to take up single or multiple bead(s) (fig. S6J-K). Compared to MdMs and monocytes, KCs were the most proficient at capturing beads in the sinusoids (fig. S6L,M).

After establishing the baseline phenotype of KC phagocytosis during MASLD, we then repeated this *in situ* phagocytosis assay with KC^Cre^ and KC^Tfeb^ mice after 16 weeks of HFHS diet (Fig. 6J). Consistent with previous data, TFEB overexpression maintained KC numbers by absolute quantification and percentage (fig. S6N). We found that TFEB did not alter the phagocytic capacity of KCs (Fig. 6K-L). However, the total number of beads captured in the liver was significantly increased in KC^Tfeb^ compared to KC^Cre^ mice (Fig. 6M). Thus, enhancing KC survival in the KC^Tfeb^ mice facilitates the uptake of beads from circulation by macrophages. As before, we observed that KCs are more efficient at ingesting beads than MoKCs and LAMs (Fig. 6M), suggesting that KC filtration cannot be fully replaced by other macrophage populations. Together these data suggest that preserving KC numbers *in vivo* can improve sinusoidal filtration.

### TFEB induction reduces oxidative stress-mediated cell death in macrophages

Having established that KC preservation is important for liver function in MASLD, we next investigated the mechanism through which TFEB induction attenuates macrophage cell death. First, we asked whether metabolic processes governed by CPT2, LAL, or GDF15 might contribute to protection of TFEB-KCs. Upon HFHS diet feeding we found that loss of CPT2, LAL, or GDF15 was dispensable for the survival of TFEB-KCs (Fig. 7A) despite effective knockdown (fig. S5G, M). This was also true in the CDAA diet for TFEB-KCs with CPT2 or GDF15 deleted (Fig. 7B, C), whereas the protective effect of TFEB in LAL KO KCs was slightly attenuated (Fig. 7C). In CPT2 and GDF15 deficient-KC^Tfeb^ mice, MdMs number was also decreased after HFHS feeding (fig. S7A-B). However, the loss of LAL in KC^Tfeb^ mice diminished the protection of TFEB against MdM accumulation (fig. S7C).

**Figure 7.**
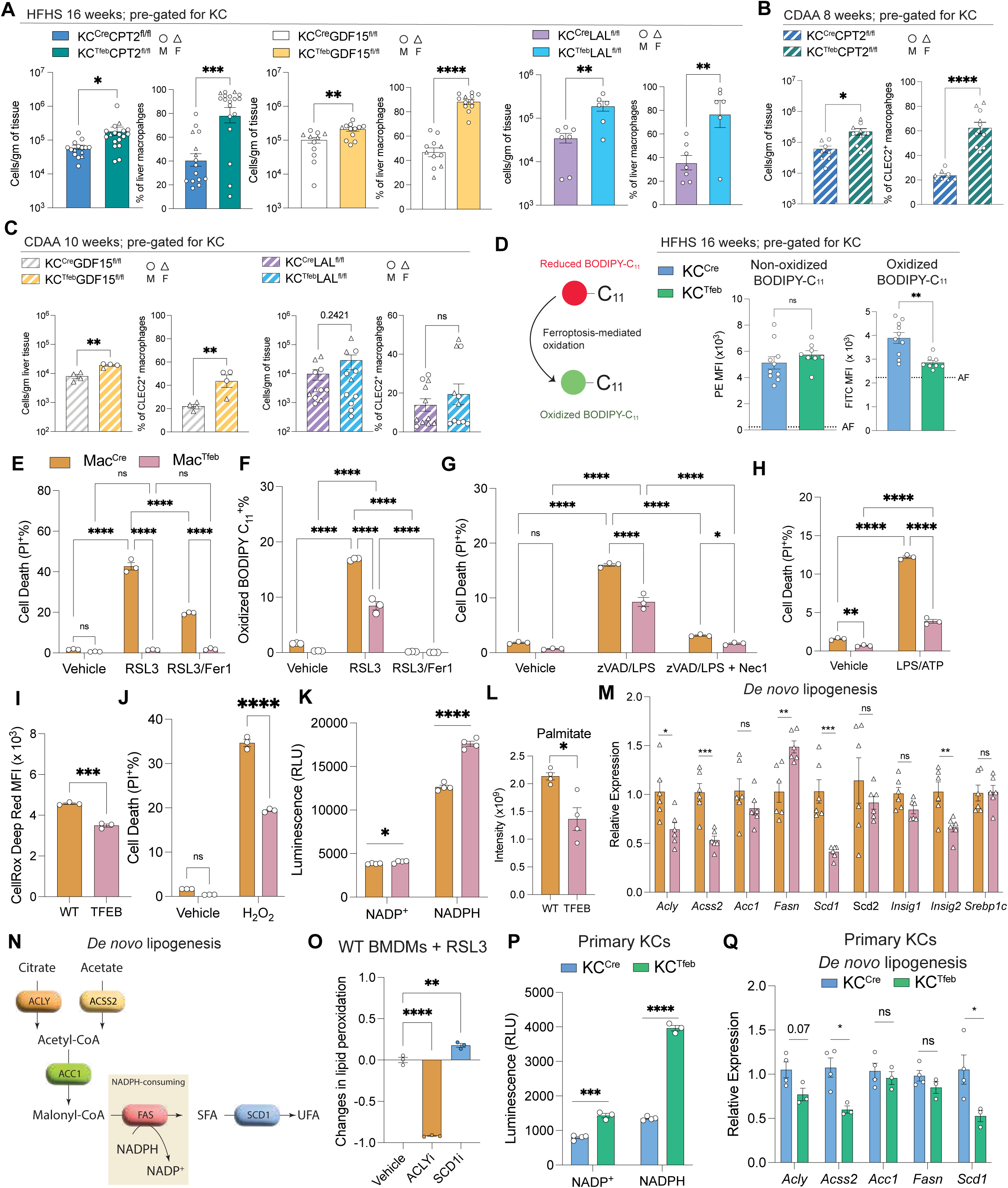
TFEB-mediated protection of macrophages *in vivo* and *in vitro*. (**A-C**) Flow cytometric quantification of KCs per gram of liver and percentages isolated from various mouse lines fed (A) HFHS diet for 16 weeks (n = 6-18/group) or (B-C) CDAA for 8-10 weeks (n = 4-12/group); circles represent males, and the triangles represent females. (**D**) MFI of BODIPY-C_11_ staining in KCs isolated from KC^Cre^ and KC^Tfeb^ mice fed HFHS for 16 weeks (n = 8-10/group). AF = autofluorescence of KCs determined from unstained samples. (**E-M)** *In vitro* experiments using BMDMs isolated from Mac^Cre^ and Mac^Tfeb^ mice. Flow cytometric percentage of (E) propidium iodide^+^ (PI^+^) signal and (F) oxidized BODIPY-C11^+^ signal in cells treated with 5µM RSL3 with/without 5µM Fer1 for 24h and 3h, respectively. Vehicle = DMSO. (G) Cells were stimulated with 20µM zVAD and 100ng/mL LPS ± 20µM necrostatin-1 (Nec1) for 24h and cell death (necroptosis) was measured by PI. Vehicle = DMSO/PBS. (H) Cells were stimulated with 500ng/mL LPS for 4h then 5mM ATP for 1h and cell death (pyroptosis) was measured by PI. Vehicle = PBS. (I) MFI of CellRox Deep Red signal in WT- or TFEB-BMDMs treated with 5µM RSL3 +/- 5µM Fer1 for 3h. Representative histograms are shown. (J) Cells were stimulated with 2.5mM H_2_O_2_ for 2h and cell death was measured by PI. (K) NADP^+^ and NADPH level in BMDMs. (L) Palmitate level measured by mass spectrometry. (M) Gene expression of DNL- related enzymes in BMDMs. (**N**) Schematic of *de novo* lipogenesis. SFA: saturated fatty acid; UFA: unsaturated fatty acid. (**O**) Change in oxidized BODIPY-C_11_ signal between inhibitor- treated and vehicle-treated WT-BMDMs 3h post-RSL3 stimulation. Cells were pretreated with respective inhibitors for 16h before co-treatment with RSL3. (**P-Q**) *Ex vivo* experiments with primary KCs isolated from isolated from male KC^Cre^ and KC^Tfeb^ mice (n = 3-4/genotype). (P) NADP^+^ and NADPH level in primary KCs. (Q) Gene expression of DNL-related enzymes in primary KCs. Data represents (A-D,P-Q) individual biological replicates or (E-L, O) technical replicates and are presented as means ±SEM. P-values were calculated using (A-D, K-M, P-Q) unpaired two-tailed Student’s t-tests, (E-J) two-way ANOVA followed by multiple t-tests, and (O) one-way ANOVA followed by multiple t-tests. NS = not significant, *p < 0.05, **p< 0.01, ***p< 0.001, ****p<0.0001.

Ferroptosis is a mode of cell death driven by lipid peroxidation that occurs secondary to iron-mediated oxidative stress(*51, 52*). This mode of death has been recently shown to contribute to KC death during MASLD(*34*). In ferroptosis, the oxidation of polyunsaturated fatty acids (PUFA) triggers loss of membrane integrity and cell death. Therefore, we assessed lipid peroxidation in KCs from WT and transgenic mice after 16-week HFHS diet feeding using BODIPY-C_11_, a lipid probe that changes fluorescence upon oxidation(*34, 53*). Although uptake of the BODIPY-C_11_ tracer was similar between WT and TFEB-KCs, lipid peroxidation was significantly reduced in TFEB-KCs (Fig. 7D). To further assess this *in vitro*, we treated WT and TFEB-BMDMs with the ferroptosis inducer RSL3. RSL3 induced cell death in WT BMDMs, but cell death was significantly attenuated in TFEB-BMDMs (Fig. 7E). As a control, we demonstrated that RSL3- induced cell death was significantly reduced by the ferroptosis inhibitor ferrostatin-1 (Fer1) (Fig. 7E). Lipid peroxidation was also attenuated in TFEB-BMDMs and completely prevented by Fer1 (Fig. 7F). A similar phenotype was observed with another inducer of ferroptosis, ML162 (fig. S7D). Time course analyses on BMDMs treated with RSL3 revealed that lipid peroxidation peaked at 6h post incubation in WT cells, whereas this was suppressed in TFEB-BMDMs at all time points (fig. S7E). Cell death of WT cells began to increase significantly 12h post-stimulation; however, TFEB BMDMs maintained their membrane integrity and circumvented cell death (fig. S7E). In addition to ferroptosis, TFEB-BMDMs were also protected from other forms of cell death including necroptosis and pyroptosis (Fig. 7G, H). Pyroptosis-mediated IL-1β release was also dampened in TFEB-BMDMs (fig. S7F). The protection against multiple death pathways suggested that TFEB acts to block common pathway in cell death. Reactive oxygen species (ROS) is a common denominator among various modes of cell death (*54–56*). We thus assessed whether intracellular ROS are reduced by TFEB induction. Utilizing the general ROS sensors CellRox, we found that TFEB-BMDMs have reduced levels of ROS compared to their WT counterpart (Fig. 7I). Furthermore, TFEB-BMDMs were resistant cell death induced by the direct ROS activator hydrogen peroxide (H_2_O_2_) (Fig. 7J). Based on our *in vivo* and *in vitro* models, TFEB reduces ROS accumulation and protects macrophages from cell death.

### TFEB promotes NADPH accumulation by suppressing de novo lipogenesis

To assess the mechanism of ROS-resistance in TFEB-BMDMs, we first evaluated the expression of genes involved in antioxidant pathways. Antioxidant genes such as those for superoxide dismutase (*Sod1/2),* glutathione peroxidase (*Gpx4*), catalase *(Cat),* and peroxidase *(Prdx1)* were decreased in TFEB-BMDMs (fig. S7G). Antioxidant pathways are fueled by reducing equivalents from NADPH(*57–59*). Therefore, we measured cellular NADP^+^ and NADPH levels in BMDMs and found that increased TFEB led to significantly higher levels of NADPH compared to WT cells (Fig. 7K). The NADPH pool can be altered via increasing production or reducing consumption. The pentose phosphate pathway (PPP) is a major pathway responsible for the generation of NADPH and its activity is sensitive to NADP^+^/NADPH level (*60–62*). The expression of genes involved in the PPP were reduced in TFEB-BMDMs suggesting that increased flux through this pathway was unlikely to explain the increase in NADPH levels (fig. S7H). Likewise, ribose-5-phosphate, the end product of oxidative PPP, was unchanged (fig. S7I). *De novo* lipogenesis (DNL) is a major NADPH-consuming pathway(*62*). We also found that palmitate, the major product of DNL, was significantly reduced in TFEB-BMDMs (Fig. 7L). Therefore, we evaluated the mRNA expression of core DNL enzymes and found that TFEB- BMDMs had significantly lower levels of *Acly*, *Acss2*, and *Scd1* (Fig. 7M). ACLY is upstream of the main NADPH consuming step of DNL whereas SCD1 is downstream (Fig. 7N). As both *Acly* and *Scd1* are reduced in TFEB-BMDMs, we evaluated the relevance of these pathways to ROS protection by treating WT cells with inhibitors of ACLY or SCD1 followed by RSL3 to induce ferroptosis. Inhibition of ACLY with BMS303141 strongly attenuated lipid peroxidation whereas SCD1 inhibition with CAY 10566 mildly exacerbates it (Fig. 7O, fig. S7J), consistent with the idea that NADPH consumption by DNL can sensitize macrophages to ferroptosis. To confirm these findings *in vivo* we quantified NADPH levels in freshly isolated primary KCs from KC^Cre^ and KC^Tfeb^ mice. We found that TFEB-overexpressing KCs had an increase in cellular NADPH (Fig. 7P). The expression of PPP-related genes was also lower than wild type (fig. S7K) and consistent with *in vitro* models, TFEB-KCs also had decreased expression of DNL-related genes (Fig. 7Q).

## DISCUSSION

The study of macrophages in MASLD pathogenesis has largely focused on recruited MdMs as drivers of tissue pathology. Metabolic liver disease is also associated with KC loss; however, the local and systemic consequences of resident macrophage attrition has not been investigated. KCs are important for liver homeostasis and immune regulation leading us to hypothesize that restoring KCs in MASLD could be an approach to attenuate liver pathology. We report here that activating TFEB in resident KCs during MASLD reprograms their metabolism, leading to reduced liver steatosis and improved KC survival.

Our data adds to a growing body of literature supporting a beneficial role for macrophage TFEB in metabolic disease. Previous studies using LysM-Cre to drive TFEB overexpression in myeloid cells revealed protection against atherosclerosis, diet-induced obesity, and insulin resistance(*43, 49*). In addition, it was recently shown that loss of Hif-2α in macrophages increases TFEB activation in KCs and reduces liver steatosis and damage(*33*). In obesity and insulin resistance, the protective effect of TFEB-overexpression in macrophages using a *Lys2-Cre* system was mediated by increased release of the anorexigenic cytokine GDF15 (*49*). While the expression of *Gdf15* mRNA was also increased in TFEB-KCs, we did not observe increased levels of GDF15 in circulation. In line with this, KC^Tfeb^ mice were not lean and had similar systemic metabolic dysfunction compared to controls. In addition, using a genetic approach to delete GDF15 from TFEB-overexpressing KCs we also found that this cytokine did not impact on liver phenotypes. Taken together, these data indicate that activating TFEB in macrophages is beneficial in obesity-related disorders through GDF15-dependent and independent mechanisms.

Several recent studies have suggested that macrophages interact with parenchymal cells to orchestrate lipid handling. This is most notable in the adipose tissue where ATMs take up and process adipocyte-derived extracellular vesicles that are rich in TAG (*63*). We recently demonstrated MdMs also take up lipids released from hepatocytes early in MASLD(*46*). In contrast, KCs isolated from steatotic livers contain fewer lipid droplets and appear less adapted to handle excess fatty acids. We found that TFEB activation in KCs enhanced their ability to handle lipids from hepatocytes and this resulted in reduced steatosis and liver injury. This conclusion is supported by the following observations:1) KC^Tfeb^ mice had reduced macrovesicular steatosis and decreased levels of several TAG species in the liver without directly altering hepatocyte lipid metabolic gene expression, 2) RNAseq of isolated KCs highlighted enhanced expression of genes involved in lipid uptake (*Cd36*) and metabolism (*Lipa*), 3) EM of TFEB-KCs revealed a significant increase in the number of lipid droplets in vacuoles compared to WT-KCs, and 4) disruption of FAO or lysosomal lipolysis in TFEB-KCs mitigated the protective effect of TFEB on hepatic steatosis. Thus, KCs are poorly adapted to the lipid-rich environment of MASLD but can be programmed by TFEB induction to enhance lipid handling and improve hepatic steatosis.

KCs that overexpress TFEB were also found to be resistant to cell death in mouse models of MASLD and MASH. Ferroptosis is a mode of cell death that occurs via iron mediated lipid peroxidation which has been implicated in KC death (*34*). We found that TFEB overexpression led to the protection of macrophages from lipid peroxidation and ferroptotic cell death. Interestingly, TFEB macrophages also had attenuated necroptosis and pyroptosis. ROS is a shared mechanism involved in these forms of cell death. Consistent with TFEB improving antioxidant defenses, cell death was also reduced in response to the ROS activator hydrogen peroxide. The observed protection against ROS was linked to increased NADPH levels in TFEB-overexpressing macrophages. Although PPP is the main NADPH-generating pathway, the expression of genes involved in this shunt was downregulated in TFEB macrophages. In contrast, DNL consumes significant NADPH and genes related to this were also downregulated in TFEB-overexpressing macrophages. Interestingly, activation of TFE3, a related family member of TFEB, has also been reported to reduce DNL in hepatocytes(*64*). Inhibiting ACLY, which blocks DNL upstream of NADPH consumption, protected WT macrophages against lipid peroxidation and ferroptosis. Thus, reprogramming of cellular metabolism via TFEB overexpression leads to altered DNL which protects macrophages from oxidative stress via increasing the intracellular stores of NADPH. We subsequently validated the reduced expression of DNL genes and enhanced levels of NADPH in KCs from TFEB overexpressing mice. Taken together our data argues that TFEB expression in KCs alters lipid biosynthesis and reduces oxidative stress, thereby improving KC fitness in MASLD.

MASLD is associated with perturbations in gut integrity and microbiota composition which influences the contents of portal blood entering the liver(*11, 65–68*). The enrichment of KCs in the periportal region and their localization to the lumen serves as a firewall to capture infiltrating bacteria/toxins and prevent systemic dissemination (*11, 13, 69, 70*). We found that preserving KC numbers through TFEB induction in the MASLD liver allowed them to maintain their fundamental role as a filter of circulation. Our data also highlighted the fact that compared to KCs, recruited macrophages such as MoKCs and LAMs fail to efficiently capture particulate antigen in the liver sinusoids of steatotic livers. This result may reflect both tissue localization and/or differences in phagocytic capacity; however, it supports that concept that recruited macrophage are unable to immediately fulfill the responsibility of the KC network. Recent data has shown that in fibrotic liver disease MdMs can form syncytia to capture bacteria when KC function is compromised (*12*), but whether this occurs in steatotic liver disease warrants future investigation.

MASLD and its sequelae remain the leading causes of liver transplant and cardiovascular death. Our data demonstrate that harnessing lysosomal lipid metabolism through TFEB induction promotes KC fitness in MASLD which leads to improved liver steatosis. To capitalize on the translational potential of TFEB will require a better understanding of the relevant pathways that are engaged downstream of this transcription factor in metabolic disease. While TFEB activators such as trehalose(*43, 71, 72*) and mTOR antagonists(*64*) are available for clinical use, more directed downstream targets would be ideal. In summary, we demonstrate that TFEB protects KCs in MASLD and this reduces liver steatosis and preserves liver filtration.

## MATERIALS AND METHODS

### Study Design

Our study generated a mouse model in which overexpression of TFEB is specific to KCs to investigate the impact of TFEB induction on KC biology and disease outcome during MASLD. We fed mice two different MASLD/MASH-inducing diets and characterized KC function and survival using flow cytometry, imaging, and lipidomics. Male mice were predominately used for obesogenic diet studies and both female and male mice were used for fibrogenic diet studies. We also generated additional depletion of lipid metabolic machinery in the KC^Tfeb^ mice to investigate the mechanism for TFEB’s influence *in vivo*. Complementary to our KC-specific induction of TFEB, BMDMs generated from pan-macrophage Cre-driven TFEB-overexpressing mice were utilized for *in vitro* studies. Sample size determination was informed by prior studies. Sample exclusion criteria (liver cancer, bite wounds, splenomegaly, and other unexpected organ abnormalities) were determined before data acquisition. Blinding was implemented whenever possible for data collection and researchers were unblinded for data analyses.

### In vivo animal studies

Wild-type male C57BL/6 animals were purchased from Jackson Laboratory. Clec4f^Cre^ and Flox/stop/flox-TFEB-3xFLAG transgenic mice were obtained from Drs. Charlotte Scott/Martin Guilliams (*16, 17*) and Dr. Andrea Ballabio(*38, 39*), respectively. KC-TdT reporter mouse were bred in-house with by crossing Clec4f^Cre^ mice with ROSA26^flox/stop/flox^-TdTomato mice. LysM^Cre^ (Mac^Cre^) and TFEB^fl/fl^ mice were obtained from Dr. Babak Razani. CPT2^fl/fl^ mice were obtained from Dr. Jennifer Philips. LAL^flox/flox^ mice were obtained from Dr. Ali Javaheri. Cx3Cr1^CreER^-ROSA26^flox-stop-flox-TdT^ (Cx3Cr1^CreER-TdT^) mice were obtained from Dr. Kory Lavine. GDF15^fl/fl^ mice were generated by CRISPR-Cas9 system to insert flox/stop/flox cassettes around the 2^nd^ exon. All mice were bred in-house at Washington University School of Medicine, St. Louis. High fat, high sucrose diet (HFHS) (42%Kcal fat diet with increased sucrose and 1.25% cholesterol) was purchased from Envigo Teklad (Cat# TD.120528). Choline-deficient, amino acid-defined diet (CDAA) was purchased from Research Diets (Cat# A06071309). All animals were fed *ad libitum* with unlimited access to drinking water. For tamoxifen injection in Cx3Cr1^CreER-TdT^ mice, 20µg tamoxifen per gram of animal was prepared fresh in corn oil by incubation at 42°C with agitation until dissolution prior to intraperitoneal (i.p.) injection. For BrdU injection, mice were injected i.p. with 100mg BrdU (resuspended in sterile PBS) per kg of animal with insulin syringes. Littermates with different genotypes were co-housed to minimize differences due to microbiota. All mice were housed in specific pathogen-free conditions. Male and female mice of 8-12 weeks of age were used for diet studies. All animal protocols were approved by the Institutional Animal Care and Use Committee at Washington University School of Medicine.

### Tissue harvest and processing

Animals were fasted for 4h before tissue harvest. Livers were collected based on our previously published protocol(*73*). Briefly, animals were euthanized by CO_2_ inhalation with cervical dislocation as secondary death confirmation. The liver was perfused with PBS through the portal vein and then dissected out. Gallbladder was discarded. Approximately 1g of the left lobe was weighed, minced, and transferred to collagenase- and DNaseI-containing (0.75mg/mL and 50ug/mL, respectively) media on ice until all the organs from all the animals were harvested. Liver in enzymatic solution was then placed on a rotating shaker for 30min at 37°C for digestion. Liver digest was then passed through 70µm cell strainer and washed with cold complete DMEM containing fetal bovine serum (FBS) to deactivate enzymatic activities. Hepatocytes were pelleted by centrifuging at 50xg, 3min, at 4°C and discarded. The supernatant was transferred to a new 50mL falcon tube and non-parenchymal cells (NPC) were pelleted at 900rpm, 7min, at 4°C. After discarding supernatant from the NPC pellet, red blood cells were lysed by resuspending cell pellets with 1mL of ACK lysis buffer (Corning) and incubated for 5min at room temperature. Ten mL of PBS was then added, and cells were re-pelleted at 900rpm, 7min, at 4°C. This cell pellet was then used for downstream analyses such as flow cytometry.

### Endolysosomal assessment, ferroptosis detection in primary cells, and in situ bead injection

Livers were perfused with PBS followed by 37°C collagenase A (1mg/mL) until tissue disintegrated. The liver was carefully dissected out and torn into smaller pieces in the collagenase-containing media, and put on a 37°C air shaker for additional digestion for 20min. After differential centrifugation described above to remove hepatocytes and pellet NPCs, NPCs were resuspended in 5mL of 35% Percoll and layered onto 5mL 70% Percoll and centrifuged for 20min at 2200 rpm at room temperature without break or acceleration. The interlayer cell suspension (enriched in macrophages) is collected into a new tube, washed with 10mL FBS-containing DMEM, and repelleted at 900 rpm for 7min at 4°C. Cells were counted and ∼4 x 10^5^ cells were incubated with 250uL of various substrates prepared in serum-containing DMEM. DQ-OVA (5µg/mL, ThermoFisher, Cat# D12053), pHrodo-red (50ug/mL, ThermoFisher, Cat#P10361), and TMR Dextran-1000MV (50µg/mL, Invitrogen, Cat#D1817) were incubated with cells for 30min at 37°C. Lysotracker green (1:2000, Invitrogen, Cat#L7526) was incubated with cells for 5min at 37°C. For *ex vivo* lipid uptake assay, BODIPY-C_16_ (1µM, ThermoFisher, Cat#D3821) was incubated for 1min at 37°C followed by quenching with 500µM of phloretin. For ferroptosis detection in KCs, NPCs were isolated based on the methods above, and incubated with 100uL of BODIPY 581/591 C_11_ (5µM, ThermoFisher, D3861) in FBS-containing DMEM for 30min at 37°C. Afterward, cells were washed with PBS and subjected to antibody staining for flow cytometry.

For *in situ* injection of fluorescent beads, the mouse was euthanized, and liver was first perfused with PBS. Then 5mL of pre-warmed, 37°C fluorescent beads (ThermoFisher, Cat#13083) prepared at 10^8^ beads/mL in PBS was slowly injected into the portal vein of the mouse over 1 min. Meanwhile, the inferior vena cava (IVC) was clamped to allow for saturation of the bead solution in the liver for 5 min. Then the IVC clamp was released, and the liver was perfused with PBS for 2min to wash off unattached beads. The liver was dissected out and incubated in 5mL of FBS-containing DMEM for 30min at 37°C. Then parts of the livers were either fixed for immunofluorescence or digested with collagenase and DNAse I following the protocol above for flow cytometry. To calculate the total beads captured per gram of tissue, beads^+^ percentages were multiplied by respective cell numbers per gram of tissue.

### Cell culture

Cells from male mice were generally used unless specified. To generate BMDMs, bone marrow was isolated from femur and tibia of mice. Briefly, muscles were removed from the bones and the ends of each bone were cut open with a razor blade. Sterile PBS was then injected into each bone to flush out the bone marrows. Marrows were pelleted at 950rpm, 7min, 4°C and resuspended in 10% CMG-conditioned DMEM (high glucose DMEM with 10% FBS, 10% CMG-conditioned media, 1% P/S, 1% L-glutamine, and 1% sodium pyruvate) for plating on bacterial culture plates for 6 days. Additional 10% CMG-conditioned DMEM was supplemented to BMDM culture on days 3 and 5. For experiments, cells were washed, trypsinized, and counted for plating in 5% CMG-conditioned DMEM on day 6 of culture. CMG14-12 cell line was expanded to confluency in complete DMEM (high glucose DMEM + 10% FBS, 1% P/S, 1% 200mM L-glutamine, and 1% 100mM sodium pyruvates) and conditioned media were collected and filtered every other day. For flow cytometry-based experiments, cells were plated on suspension plates at 5 x 10^5^ cells/mL; for all other assays, cells were plated on tissue culture-treated plates at the same density.

For primary KC culture to obtain RNA and NADP^+^/NADPH assay, livers were perfused with collagenase as described earlier to maximize yield. After differential centrifugation, CD45^+^ cells were positively selected using CD45 microbeads and MS columns (Miltenyi Biotec, Cat# 130-052-301; #130-042-201) following manufacturer’s protocol. CD45^+^ cells from one mouse were plated onto tissue culture plates for 2h incubation at 37°C to allow adhesion. Non-adherent cells were then washed off with PBS twice, and cells were lysed for RNA.

## Acknowledgments

We thank K. Lavine and G. Randolph for helpful advice and reagents. We also thank the Tissue Analysis Core of the Digestive Disease Research Core Center for histological staining, Genome Technology Access Center at the McDonnell Genome Institute for assistance with genomic analysis, Metabolomics Facility for lipidomic studies, Metabolic Tissue Function Core for Seahorse equipment, and Flow Cytometry & Fluorescence Activated Cell Sorting Core for equipment at Washington University School of Medicine. The Genome Technology Access Center is partially supported by NCI Cancer Center Support Grant #P30 CA91842 to the Siteman Cancer Center. This publication is solely the responsibility of the authors and does not necessarily represent the official view of NCRR or NIH. The authors have no conflicting financial interests.

## Data and materials availability

All data are available upon request from the corresponding authors.

## Author contributions

Conceptualization: JDS, BR, MMC; Methodology: MMC, JDS, SD; Investigation: MMC, SD, WB, KB, NF, ZG, AP, CFF, KF, LH, BQY, CP; Visualization: MMC, WB; Funding acquisition: JDS, BR; Project administration: JDS; Supervision: JDS; Writing – original draft: MMC, JDS; Writing – review & editing: MMC, JDS, SD, WB, KB, NF, ZG, AP, CFF, KF, LH, BQY, CP, AJ

## List of Abbreviations

KC: Kupffer cells
TFEB: Transcription factor EB
MASLD: metabolic dysfunction-associated steatotic liver diseases
MdM: monocyte-derived macrophages
MASH: metabolic dysfunction-associated steatohepatitis
GDF15: growth differentiation factor 15
T2D: type 2 diabetes
scRNAseq: single-cell RNA sequencing
FA: fatty acid
BMDM: bone marrow-derived macrophages
TdT: TdTomato
HFHS: high fat, high sucrose and high cholesterol (diet)
STD: standard diet
MoKC: monocyte-derived KC
LAM: lipid-associated macrophages
CDAA: choline-deficient amino acid-defined (diet)
TAG: triglyceride
ALT: alanine transaminase
FAO: fatty acid oxidation
DEG: differentially expressed genes
ROS: reactive oxygen species
DNL: *de novo* lipogenesis
PI: propidium iodide

## References and Note

1. D. G. Tiniakos, M. B. Vos, E. M. Brunt, Nonalcoholic Fatty Liver Disease: Pathology and Pathogenesis. Annual Review of Pathology: Mechanisms of Disease 5, 145–171 (2010).

2. P. S. Dulai, S. Singh, J. Patel, M. Soni, L. J. Prokop, Z. Younossi, G. Sebastiani, M. Ekstedt, H. Hagstrom, P. Nasr, P. Stal, V. W. Wong, S. Kechagias, R. Hultcrantz, R. Loomba, Increased risk of mortality by fibrosis stage in nonalcoholic fatty liver disease: Systematic review and meta-analysis. Hepatology 65, 1557–1565 (2017).

3. Z. M. Younossi, D. Blissett, R. Blissett, L. Henry, M. Stepanova, Y. Younossi, A. Racila, S. Hunt, R. Beckerman, The economic and clinical burden of nonalcoholic fatty liver disease in the United States and Europe. Hepatology 64, 1577–1586 (2016).

4. Z. Younossi, F. Tacke, M. Arrese, B. Chander Sharma, I. Mostafa, E. Bugianesi, V. Wai-Sun Wong, Y. Yilmaz, J. George, J. Fan, M. B. Vos, Global Perspectives on Nonalcoholic Fatty Liver Disease and Nonalcoholic Steatohepatitis. Hepatology 69, 2672–2682 (2019).

5. G. Targher, C. D. Byrne, Nonalcoholic Fatty Liver Disease: A Novel Cardiometabolic Risk Factor for Type 2 Diabetes and Its Complications. The Journal of Clinical Endocrinology & Metabolism 98, 483–495 (2013).

6. S. A. Harrison, R. Taub, G. W. Neff, K. J. Lucas, D. Labriola, S. E. Moussa, N. Alkhouri, M. R. Bashir, Resmetirom for nonalcoholic fatty liver disease: a randomized, double-blind, placebo-controlled phase 3 trial. Nat Med 29, 2919–2928 (2023).

7. S. A. Harrison, P. Bedossa, C. D. Guy, J. M. Schattenberg, R. Loomba, R. Taub, D. Labriola, S. E. Moussa, G. W. Neff, M. E. Rinella, Q. M. Anstee, M. F. Abdelmalek, Z. Younossi, S. J. Baum, S. Francque, M. R. Charlton, P. N. Newsome, N. Lanthier, I. Schiefke, A. Mangia, J. M. Pericas, R. Patil, A. J. Sanyal, M. Noureddin, M. B. Bansal, N. Alkhouri, L. Castera, M. Rudraraju, V. Ratziu, M.-N. Investigators, A Phase 3, Randomized, Controlled Trial of Resmetirom in NASH with Liver Fibrosis. N Engl J Med 390, 497–509 (2024).

8. E. Barreby, P. Chen, M. Aouadi, Macrophage functional diversity in NAFLD - more than inflammation. Nat Rev Endocrinol 18, 461–472 (2022).

9. Z. Liu, Y. Gu, S. Chakarov, C. Bleriot, I. Kwok, X. Chen, A. Shin, W. Huang, R. J. Dress, C. A. Dutertre, A. Schlitzer, J. Chen, L. G. Ng, H. Wang, Z. Liu, B. Su, F. Ginhoux, Fate Mapping via Ms4a3-Expression History Traces Monocyte-Derived Cells. Cell 178, 1509–1525 e1519 (2019).

10. B. A. David, R. M. Rezende, M. M. Antunes, M. M. Santos, M. A. Freitas Lopes, A. B. Diniz, R. V. Sousa Pereira, S. C. Marchesi, D. M. Alvarenga, B. N. Nakagaki, A. M. Araujo, D. S. Dos Reis, R. M. Rocha, P. E. Marques, W. Y. Lee, J. Deniset, P. X. Liew, S. Rubino, L. Cox, V. Pinho, T. M. Cunha, G. R. Fernandes, A. G. Oliveira, M. M. Teixeira, P. Kubes, G. B. Menezes, Combination of Mass Cytometry and Imaging Analysis Reveals Origin, Location, and Functional Repopulation of Liver Myeloid Cells in Mice. Gastroenterology 151, 1176–1191 (2016).

11. Y.-H. Han, E. J. Onufer, L.-H. Huang, R. W. Sprung, W. S. Davidson, R. S. Czepielewski, M. Wohltmann, M. G. Sorci-Thomas, B. W. Warner, G. J. Randolph, Enterically derived high-density lipoprotein restrains liver injury through the portal vein. Science 373, eabe6729 (2021).

12. M. Peiseler, B. Araujo David, J. Zindel, B. G. J. Surewaard, W. Y. Lee, F. Heymann, Y. Nusse, F. V. S. Castanheira, R. Shim, A. Guillot, A. Bruneau, J. Atif, C. Perciani, C. Ohland, P. Ganguli Mukherjee, A. Niehrs, R. Thuenauer, M. Altfeld, M. Amrein, Z. Liu, P. M. K. Gordon, K. McCoy, J. Deniset, S. MacParland, F. Ginhoux, F. Tacke, P. Kubes, Kupffer cell-like syncytia replenish resident macrophage function in the fibrotic liver. Science 381, eabq5202 (2023).

13. M. L. Balmer, E. Slack, A. de Gottardi, M. A. E. Lawson, S. Hapfelmeier, L. Miele, A. Grieco, H. Van Vlierberghe, R. Fahrner, N. Patuto, C. Bernsmeier, F. Ronchi, M. Wyss, D. Stroka, N. Dickgreber, M. H. Heim, K. D. McCoy, A. J. Macpherson, The Liver May Act as a Firewall Mediating Mutualism Between the Host and Its Gut Commensal Microbiota. Science Translational Medicine 6, 237ra266-237ra266 (2014).

14. A. Gola, M. G. Dorrington, E. Speranza, C. Sala, R. M. Shih, A. J. Radtke, H. S. Wong, A. P. Baptista, J. M. Hernandez, G. Castellani, I. D. C. Fraser, R. N. Germain, Commensal-driven immune zonation of the liver promotes host defence. Nature, (2020).

15. Y. Kanamori, M. Tanaka, M. Itoh, K. Ochi, A. Ito, I. Hidaka, I. Sakaida, Y. Ogawa, T. Suganami, Iron-Rich Kupffer Cells Exhibit Phenotypic Changes during the Development of Liver Fibrosis in NASH. iScience, 102032 (2021).

16. C. L. Scott, F. Zheng, P. De Baetselier, L. Martens, Y. Saeys, S. De Prijck, S. Lippens, C. Abels, S. Schoonooghe, G. Raes, N. Devoogdt, B. N. Lambrecht, A. Beschin, M. Guilliams, Bone marrow-derived monocytes give rise to self-renewing and fully differentiated Kupffer cells. Nature Communications 7, (2016).

17. C. L. Scott, W. T’Jonck, L. Martens, H. Todorov, D. Sichien, B. Soen, J. Bonnardel, S. De Prijck, N. Vandamme, R. Cannoodt, W. Saelens, B. Vanneste, W. Toussaint, P. De Bleser, N. Takahashi, P. Vandenabeele, S. Henri, C. Pridans, D. A. Hume, B. N. Lambrecht, P. De Baetselier, S. W. F. Milling, J. A. Van Ginderachter, B. Malissen, G. Berx, A. Beschin, Y. Saeys, M. Guilliams, The Transcription Factor ZEB2 Is Required to Maintain the Tissue-Specific Identities of Macrophages. Immunity 49, 312–325.e315 (2018).

18. J. Bonnardel, W. T’Jonck, D. Gaublomme, R. Browaeys, C. L. Scott, L. Martens, B. Vanneste, S. De Prijck, S. A. Nedospasov, A. Kremer, E. Van Hamme, P. Borghgraef, W. Toussaint, P. De Bleser, I. Mannaerts, A. Beschin, L. A. van Grunsven, B. N. Lambrecht, T. Taghon, S. Lippens, D. Elewaut, Y. Saeys, M. Guilliams, Stellate Cells, Hepatocytes, and Endothelial Cells Imprint the Kupffer Cell Identity on Monocytes Colonizing the Liver Macrophage Niche. Immunity, (2019).

19. K. Miura, L. Yang, N. v. Rooijen, H. Ohnishi, E. Seki, Hepatic recruitment of macrophages promotes nonalcoholic steatohepatitis through CCR2. American Journal of Physiology-Gastrointestinal and Liver Physiology 302, G1310–G1321 (2012).

20. L. Beattie, A. Sawtell, J. Mann, T. C. M. Frame, B. Teal, F. de Labastida Rivera, N. Brown, K. Walwyn-Brown, J. W. J. Moore, S. MacDonald, E.-K. Lim, J. E. Dalton, C. R. Engwerda, K. P. MacDonald, P. M. Kaye, Bone marrow-derived and resident liver macrophages display unique transcriptomic signatures but similar biological functions. Journal of Hepatology 65, 758–768 (2016).

21. M. Guilliams, J. Bonnardel, B. Haest, B. Vanderborght, C. Wagner, A. Remmerie, A. Bujko, L. Martens, T. Thoné, R. Browaeys, F. F. De Ponti, B. Vanneste, C. Zwicker, F. R. Svedberg, T. Vanhalewyn, A. Gonçalves, S. Lippens, B. Devriendt, E. Cox, G. Ferrero, V. Wittamer, A. Willaert, S. J. F. Kaptein, J. Neyts, K. Dallmeier, P. Geldhof, S. Casaert, B. Deplancke, P. ten Dijke, A. Hoorens, A. Vanlander, F. Berrevoet, Y. Van Nieuwenhove, Y. Saeys, W. Saelens, H. Van Vlierberghe, L. Devisscher, C. L. Scott, Spatial proteogenomics reveals distinct and evolutionarily conserved hepatic macrophage niches. Cell 185, 379–396.e338 (2022).

22. P. Ramachandran, R. Dobie, J. R. Wilson-Kanamori, E. F. Dora, B. E. P. Henderson, N. T. Luu, J. R. Portman, K. P. Matchett, M. Brice, J. A. Marwick, R. S. Taylor, M. Efremova, R. Vento-Tormo, N. O. Carragher, T. J. Kendall, J. A. Fallowfield, E. M. Harrison, D. J. Mole, S. J. Wigmore, P. N. Newsome, C. J. Weston, J. P. Iredale, F. Tacke, J. W. Pollard, C. P. Ponting, J. C. Marioni, S. A. Teichmann, N. C. Henderson, Resolving the fibrotic niche of human liver cirrhosis at single-cell level. Nature 575, 512–518 (2019).

23. R. G. Fred, J. Steen Pedersen, J. J. Thompson, J. Lee, P. N. Timshel, S. Stender, M. Opseth Rygg, L. L. Gluud, V. Bjerregaard Kristiansen, F. Bendtsen, T. Hansen, T. H. Pers, Single-cell transcriptome and cell type-specific molecular pathways of human non-alcoholic steatohepatitis. Scientific Reports 12, 13484 (2022).

24. S. A. MacParland, J. C. Liu, X. Z. Ma, B. T. Innes, A. M. Bartczak, B. K. Gage, J. Manuel, N. Khuu, J. Echeverri, I. Linares, R. Gupta, M. L. Cheng, L. Y. Liu, D. Camat, S. W. Chung, R. K. Seliga, Z. Shao, E. Lee, S. Ogawa, M. Ogawa, M. D. Wilson, J. E. Fish, M. Selzner, A. Ghanekar, D. Grant, P. Greig, G. Sapisochin, N. Selzner, N. Winegarden, O. Adeyi, G. Keller, G. D. Bader, I. D. McGilvray, Single cell RNA sequencing of human liver reveals distinct intrahepatic macrophage populations. Nat Commun 9, 4383 (2018).

25. C. Blériot, E. Barreby, G. Dunsmore, R. Ballaire, S. Chakarov, X. Ficht, G. De Simone, F. Andreata, V. Fumagalli, W. Guo, G. Wan, G. Gessain, A. Khalilnezhad, X. M. Zhang, N. Ang, P. Chen, C. Morgantini, V. Azzimato, W. T. Kong, Z. Liu, R. Pai, J. Lum, F. Shihui, I. Low, C. Xu, B. Malleret, M. F. M. Kairi, A. Balachander, O. Cexus, A. Larbi, B. Lee, E. W. Newell, L. G. Ng, W. W. Phoo, R. M. Sobota, A. Sharma, S. W. Howland, J. Chen, M. Bajenoff, L. Yvan-Charvet, N. Venteclef, M. Iannacone, M. Aouadi, F. Ginhoux, A subset of Kupffer cells regulates metabolism through the expression of CD36. Immunity, (2021).

26. S. Daemen, A. Gainullina, G. Kalugotla, L. He, M. M. Chan, J. W. Beals, K. H. Liss, S. Klein, A. E. Feldstein, B. N. Finck, M. N. Artyomov, J. D. Schilling, Dynamic Shifts in the Composition of Resident and Recruited Macrophages Influence Tissue Remodeling in NASH. Cell Reports 34, (2021).

27. A. Remmerie, L. Martens, T. Thoné, A. Castoldi, R. Seurinck, B. Pavie, J. Roels, B. Vanneste, S. De Prijck, M. Vanhockerhout, M. Binte Abdul Latib, L. Devisscher, A. Hoorens, J. Bonnardel, N. Vandamme, A. Kremer, P. Borghgraef, H. Van Vlierberghe, S. Lippens, E. Pearce, Y. Saeys, C. L. Scott, Osteopontin Expression Identifies a Subset of Recruited Macrophages Distinct from Kupffer Cells in the Fatty Liver. Immunity, (2020).

28. S. Tran, I. Baba, L. Poupel, S. Dussaud, M. Moreau, A. Gélineau, G. Marcelin, E. Magréau-Davy, M. Ouhachi, P. Lesnik, A. Boissonnas, W. Le Goff, B. E. Clausen, L. Yvan-Charvet, F. Sennlaub, T. Huby, E. L. Gautier, Impaired Kupffer Cell Self-Renewal Alters the Liver Response to Lipid Overload during Non-alcoholic Steatohepatitis. Immunity, (2020).

29. J. S. Seidman, T. D. Troutman, M. Sakai, A. Gola, N. J. Spann, H. Bennett, C. M. Bruni, Z. Ouyang, R. Z. Li, X. Sun, B. T. Vu, M. P. Pasillas, K. M. Ego, D. Gosselin, V. M. Link, L.-W. Chong, R. M. Evans, B. M. Thompson, J. G. McDonald, M. Hosseini, J. L. Witztum, R. N. Germain, C. K. Glass, Niche-Specific Reprogramming of Epigenetic Landscapes Drives Myeloid Cell Diversity in Nonalcoholic Steatohepatitis. Immunity 52, 1057–1074.e1057 (2020).

30. G. N. Ioannou, C. S. Landis, G. Y. Jin, W. G. Haigh, G. C. Farrell, R. Kuver, S. P. Lee, C. Savard, Cholesterol Crystals in Hepatocyte Lipid Droplets Are Strongly Associated With Human Nonalcoholic Steatohepatitis. Hepatol Commun 3, 776–791 (2019).

31. G. N. Ioannou, W. G. Haigh, D. Thorning, C. Savard, Hepatic cholesterol crystals and crown-like structures distinguish NASH from simple steatosis. J Lipid Res 54, 1326–1334 (2013).

32. Z. Lei, J. Yu, Y. Wu, J. Shen, S. Lin, W. Xue, C. Mao, R. Tang, H. Sun, X. Qi, X. Wang, L. Xu, C. Wei, X. Wang, H. Chen, P. Hao, W. Yin, J. Zhu, Y. Li, Y. Wu, S. Liu, H. Liang, X. Chen, C. Su, S. Zhou, CD1d protects against hepatocyte apoptosis in non-alcoholic steatohepatitis. J Hepatol 80, 194–208 (2024).

33. I. Jeelani, J.-S. Moon, F. F. da Cunha, C. A. Nasamran, S. Jeon, X. Zhang, G. K. Bandyopadhyay, K. Dobaczewska, Z. Mikulski, M. Hosseini, X. Liu, T. Kisseleva, D. A. Brenner, S. Singh, R. Loomba, M. Kim, Y. S. Lee, HIF-2α drives hepatic Kupffer cell death and proinflammatory recruited macrophage activation in nonalcoholic steatohepatitis. Science Translational Medicine 16, eadi0284 (2024).

34. J. Zhang, Y. Wang, M. Fan, Y. Guan, W. Zhang, F. Huang, Z. Zhang, X. Li, B. Yuan, W. Liu, M. Geng, X. Li, J. Xu, C. Jiang, W. Zhao, F. Ye, W. Zhu, L. Meng, S. Lu, R. Holmdahl, Reactive oxygen species regulation by NCF1 governs ferroptosis susceptibility of Kupffer cells to MASH. Cell Metab 36, 1745–1763 e1746 (2024).

35. O. A. Brady, J. A. Martina, R. Puertollano, Emerging roles for TFEB in the immune response and inflammation. Autophagy 14, 181–189 (2018).

36. J. E. Irazoqui. (Elsevier Ltd, 2020), vol. 41, pp. 157–171.

37. J. A. Martina, H. I. Diab, H. Li, R. Puertollano. (Birkhauser Verlag AG, 2014), vol. 71, pp. 2483–2497.

38. C. Settembre, R. De Cegli, G. Mansueto, P. K. Saha, F. Vetrini, O. Visvikis, T. Huynh, A. Carissimo, D. Palmer, T. Jürgen Klisch, A. C. Wollenberg, D. Di Bernardo, L. Chan, J. E. Irazoqui, A. Ballabio, TFEB controls cellular lipid metabolism through a starvation-induced autoregulatory loop. Nature Cell Biology 15, 647–658 (2013).

39. C. Settembre, C. Di Malta, V. A. Polito, M. G. Arencibia, F. Vetrini, S. Erdin, S. U. Erdin, T. Huynh, D. Medina, P. Colella, M. Sardiello, D. C. Rubinsztein, A. Ballabio, TFEB Links Autophagy to Lysosomal Biogenesis. Science 332, 1429–1433 (2011).

40. R. Emanuel, I. Sergin, S. Bhattacharya, J. N. Turner, S. Epelman, C. Settembre, A. Diwan, A. Ballabio, B. Razani, Induction of Lysosomal Biogenesis in Atherosclerotic Macrophages Can Rescue Lipid-Induced Lysosomal Dysfunction and Downstream Sequelae. Arteriosclerosis, Thrombosis, and Vascular Biology 34, 1942–1952 (2014).

41. H. Martini-Stoica, Y. Xu, A. Ballabio, H. Zheng, The Autophagy–Lysosomal Pathway in Neurodegeneration: A TFEB Perspective. Trends in Neurosciences 39, 221–234 (2016).

42. Y. Liu, X. Xue, H. Zhang, X. Che, J. Luo, P. Wang, J. Xu, Z. Xing, L. Yuan, Y. Liu, X. Fu, D. Su, S. Sun, H. Zhang, C. Wu, J. Yang, Neuronal-targeted TFEB rescues dysfunction of the autophagy-lysosomal pathway and alleviates ischemic injury in permanent cerebral ischemia. Autophagy 15, 493–509 (2019).

43. T. D. Evans, S. J. Jeong, X. Zhang, I. Sergin, B. Razani. (Taylor and Francis Inc., 2018), vol. 14, pp. 724–726.

44. I. Sergin, T. D. Evans, X. Zhang, S. Bhattacharya, C. J. Stokes, E. Song, S. Ali, B. Dehestani, K. B. Holloway, P. S. Micevych, A. Javaheri, J. R. Crowley, A. Ballabio, J. D. Schilling, S. Epelman, C. C. Weihl, A. Diwan, D. Fan, M. A. Zayed, B. Razani, Exploiting macrophage autophagy-lysosomal biogenesis as a therapy for atherosclerosis. Nature Communications 8, 15750–15750 (2017).

45. J. D. Schilling, H. M. Machkovech, L. He, A. Diwan, J. E. Schaffer, TLR4 Activation Under Lipotoxic Conditions Leads to Synergistic Macrophage Cell Death through a TRIF-Dependent Pathway. The Journal of Immunology 190, 1285–1296 (2013).

46. M. M. Chan, S. Daemen, J. W. Beals, M. Terekhova, B. Q. Yang, C. F. Fu, L. He, A. C. Park, G. I. Smith, B. Razani, K. Byrnes, W. L. Beatty, S. R. Eckhouse, J. C. Eagon, D. Ferguson, B. N. Finck, S. Klein, M. N. Artyomov, J. D. Schilling, Steatosis drives monocyte-derived macrophage accumulation in human metabolic dysfunction-associated fatty liver disease. JHEP Rep 5, 100877 (2023).

47. A. Carracedo, L. C. Cantley, P. P. Pandolfi, Cancer metabolism: fatty acid oxidation in the limelight. Nat Rev Cancer 13, 227–232 (2013).

48. S. Zhang, X. Peng, S. Yang, X. Li, M. Huang, S. Wei, J. Liu, G. He, H. Zheng, L. Yang, H. Li, Q. Fan, The regulation, function, and role of lipophagy, a form of selective autophagy, in metabolic disorders. Cell Death Dis 13, 132 (2022).

49. J. Kim, S. H. Kim, H. Kang, S. Lee, S.-Y. Park, Y. Cho, Y.-M. Lim, J. W. Ahn, Y.-H. Kim, S. Chung, C. S. Choi, Y. J. Jang, H. S. Park, Y. Heo, K. H. Kim, M.-S. Lee, TFEB–GDF15 axis protects against obesity and insulin resistance as a lysosomal stress response. Nature Metabolism 3, 410–427 (2021).

50. T. Tran, J. Yang, J. Gardner, Y. Xiong, GDF15 deficiency promotes high fat diet-induced obesity in mice. PLoS One 13, e0201584 (2018).

51. S. J. Dixon, K. M. Lemberg, M. R. Lamprecht, R. Skouta, E. M. Zaitsev, C. E. Gleason, D. N. Patel, A. J. Bauer, A. M. Cantley, W. S. Yang, B. Morrison, 3rd, B. R. Stockwell, Ferroptosis: an iron-dependent form of nonapoptotic cell death. Cell 149, 1060–1072 (2012).

52. B. R. Stockwell, Ferroptosis turns 10: Emerging mechanisms, physiological functions, and therapeutic applications. Cell 185, 2401–2421 (2022).

53. A. M. Martinez, A. Kim, W. S. Yang, in Immune Mediators in Cancer: Methods and Protocols, I. Vancurova, Y. Zhu, Eds. (Springer US, New York, NY, 2020), pp. 125-130.

54. S. J. Dixon, B. R. Stockwell, The role of iron and reactive oxygen species in cell death. Nat Chem Biol 10, 9–17 (2014).

55. Y. Wang, P. Shi, Q. Chen, Z. Huang, D. Zou, J. Zhang, X. Gao, Z. Lin, Mitochondrial ROS promote macrophage pyroptosis by inducing GSDMD oxidation. J Mol Cell Biol 11, 1069–1082 (2019).

56. G. E. Villalpando-Rodriguez, S. B. Gibson, Reactive Oxygen Species (ROS) Regulates Different Types of Cell Death by Acting as a Rheostat. Oxid Med Cell Longev 2021, 9912436 (2021).

57. S. C. Lu, Regulation of glutathione synthesis. Mol Aspects Med 30, 42–59 (2009).

58. T. Sandalova, L. Zhong, Y. Lindqvist, A. Holmgren, G. Schneider, Three-dimensional structure of a mammalian thioredoxin reductase: implications for mechanism and evolution of a selenocysteine-dependent enzyme. Proc Natl Acad Sci U S A 98, 9533–9538 (2001).

59. H. N. Kirkman, G. F. Gaetani, Catalase: a tetrameric enzyme with four tightly bound molecules of NADPH. Proc Natl Acad Sci U S A 81, 4343–4347 (1984).

60. T. TeSlaa, M. Ralser, J. Fan, J. D. Rabinowitz, The pentose phosphate pathway in health and disease. Nat Metab 5, 1275–1289 (2023).

61. L. Chen, Z. Zhang, A. Hoshino, H. D. Zheng, M. Morley, Z. Arany, J. D. Rabinowitz, NADPH production by the oxidative pentose-phosphate pathway supports folate metabolism. Nat Metab 1, 404–415 (2019).

62. J. Fan, J. Ye, J. J. Kamphorst, T. Shlomi, C. B. Thompson, J. D. Rabinowitz, Quantitative flux analysis reveals folate-dependent NADPH production. Nature 510, 298–302 (2014).

63. S. E. Flaherty, 3rd, A. Grijalva, X. Xu, E. Ables, A. Nomani, A. W. Ferrante, Jr., A lipase-independent pathway of lipid release and immune modulation by adipocytes. Science 363, 989–993 (2019).

64. B. S. Gosis, S. Wada, C. Thorsheim, K. Li, S. Jung, J. H. Rhoades, Y. Yang, J. Brandimarto, L. Li, K. Uehara, C. Jang, M. Lanza, N. B. Sanford, M. R. Bornstein, S. Jeong, P. M. Titchenell, S. B. Biddinger, Z. Arany, Inhibition of nonalcoholic fatty liver disease in mice by selective inhibition of mTORC1. Science 376, eabf8271 (2022).

65. A. Nier, Y. Huber, C. Labenz, M. Michel, I. Bergheim, J. M. Schattenberg, Adipokines and Endotoxemia Correlate with Hepatic Steatosis in Non-Alcoholic Fatty Liver Disease (NAFLD). Nutrients 12, (2020).

66. L. Miele, V. Valenza, G. La Torre, M. Montalto, G. Cammarota, R. Ricci, R. Masciana, A. Forgione, M. L. Gabrieli, G. Perotti, F. M. Vecchio, G. Rapaccini, G. Gasbarrini, C. P. Day, A. Grieco, Increased intestinal permeability and tight junction alterations in nonalcoholic fatty liver disease. Hepatology 49, 1877–1887 (2009).

67. G. Carpino, M. Del Ben, D. Pastori, R. Carnevale, F. Baratta, D. Overi, H. Francis, V. Cardinale, P. Onori, S. Safarikia, V. Cammisotto, D. Alvaro, G. Svegliati-Baroni, F. Angelico, E. Gaudio, F. Violi, Increased Liver Localization of Lipopolysaccharides in Human and Experimental NAFLD. Hepatology 72, 470–485 (2020).

68. B. Araujo David, J. Atif, E. S. C. F. Vargas, T. Yasmin, A. Guillot, Y. Ait Ahmed, M. Peiseler, J. W. Hommes, L. Salm, M. A. Brundler, B. G. J. Surewaard, W. Elhenawy, S. MacParland, F. Ginhoux, K. McCoy, P. Kubes, Kupffer cell reverse migration into the liver sinusoids mitigates neonatal sepsis and meningitis. Sci Immunol 9, eadq9704 (2024).

69. A. Gola, M. G. Dorrington, E. Speranza, C. Sala, R. M. Shih, A. J. Radtke, H. S. Wong, A. P. Baptista, J. M. Hernandez, G. Castellani, I. D. C. Fraser, R. N. Germain, Commensal-driven immune zonation of the liver promotes host defence. Nature 589, 131–136 (2021).

70. J. Todoric, G. Di Caro, S. Reibe, D. C. Henstridge, C. R. Green, A. Vrbanac, F. Ceteci, C. Conche, R. McNulty, S. Shalapour, K. Taniguchi, P. J. Meikle, J. D. Watrous, R. Moranchel, M. Najhawan, M. Jain, X. Liu, T. Kisseleva, M. T. Diaz-Meco, J. Moscat, R. Knight, F. R. Greten, L. F. Lau, C. M. Metallo, M. A. Febbraio, M. Karin, Fructose stimulated de novo lipogenesis is promoted by inflammation. Nature Metabolism 2, 1034–1045 (2020).

71. S. J. Jeong, J. Stitham, T. D. Evans, X. Zhang, A. Rodriguez-Velez, Y. S. Yeh, J. Tao, K. Takabatake, S. Epelman, I. J. Lodhi, J. D. Schilling, B. J. DeBosch, A. Diwan, B. Razani, Trehalose causes low-grade lysosomal stress to activate TFEB and the autophagy-lysosome biogenesis response. Autophagy 17, 3740–3752 (2021).

72. B. J. DeBosch, M. R. Heitmeier, A. L. Mayer, C. B. Higgins, J. R. Crowley, T. E. Kraft, M. Chi, E. P. Newberry, Z. Chen, B. N. Finck, N. O. Davidson, K. E. Yarasheski, P. W. Hruz, K. H. Moley, Trehalose inhibits solute carrier 2A (SLC2A) proteins to induce autophagy and prevent hepatic steatosis. Sci Signal 9, ra21 (2016).

73. S. Daemen, M. M. Chan, J. D. Schilling, Comprehensive analysis of liver macrophage composition by flow cytometry and immunofluorescence in murine NASH. STAR Protocols 2, 100511 (2021).

