## Supplemental Information for "Induction of TFEB promotes Kupffer cell survival and reduces lipid accumulation and inflammation in MASLD"

**Supplementary Materials**

Supplemental Materials and Methods

Fig. S1 to S7

Supplemental Table 1

**Supplemental Materials and Methods**

*Flow cytometry and sorting (FACS)*

Cells were resuspended in 1:250 ZombieAqua BV510 (in PBS) and incubated on ice for 15min in the dark. After washing with FACS buffer (PBS + 2mM EDTA + 0.5% BSA w/v), cells were re-pelleted and incubated with 1:10 Fc Block (in FACS buffer) for 5min on ice in the dark. Then 90uL of antibody cocktail was added to the cells with Fc Block for 30-60min incubation on ice in the dark. For intracellular staining with anti-Ki67 (BioLegend) or anti-BrdU (BD Biosciences Cat#552598), cells were permeabilized and stained following the manufacturer’s protocol. Cells were then washed with FACS buffer and resuspended in 300-500uL FACS buffer for data acquisition using a BD Fortessa X20 flow cytometer. For cell sorting, a BD FACSAriaII was utilized to collect KCs (singlet, live, CD45^+^F480^hi^CD11b^int^TIM4^+^VSIG4^+^). Data analyses were performed using FlowJo software.

*Electron Microscopy*

KCs were FACS-purified based on TIM4 expression as described above. For ultrastructural analyses, cells were fixed in 2% paraformaldehyde/2.5% glutaraldehyde (Ted Pella Inc., Redding, CA) in 100mM sodium cacodylate buffer for 2h at room temperature. Samples were washed in sodium cacodylate buffer and postfixed in 1% osmium tetroxide (Ted Pella Inc.) for 1h at room temperature. Samples were then rinsed extensively in dH20 prior to en bloc staining with 1% aqueous uranyl acetate (Electron Microscopy Sciences, Hatfield, PA) for 1h. Following several rinses in dH20, samples were dehydrated in a graded series of ethanol, and embedded in Eponate 12 resin (Ted Pella Inc). Ultrathin sections of 95nm were cut with a Leica Ultracut UCT ultramicrotome (Leica Microsystems Inc., Bannockburn, IL), stained with uranyl acetate and lead citrate, and viewed on a JEOL 1200 EX transmission electron microscope (JEOL USA Inc., Peabody, MA) equipped with an AMT 8 megapixel digital camera and AMT Image Capture Engine V602 software (Advanced Microscopy Techniques, Woburn, MA). For quantitation of lipid droplets, 20 to 40 cells that had a nucleus cut in cross-section (indicating cross-section through the middle of cell) were randomly chosen, and images of each cell were taken at a magnification of 5,000X. The cross-sectional area of each of the lipid droplets and the cytosol of corresponding cell were determined using Image J 1.38g (National Institutes of Health, USA, customized for AMT images). Data is expressed as total number of lipid droplets for each cell type, and the total cross-sectional area of lipid droplets per total area of cytosol for each cell type. Distribution histogram was generated using R ggplot2 package.

*Bulk RNA sequencing*

Total RNA integrity was determined using Agilent Bioanalyzer or 4200 Tapestation. Library preparation was performed with 10ng of total RNA with a Bioanalyzer RIN score greater than 8.0. ds-cDNA was prepared using the SMARTer Ultra Low RNA kit for Illumina Sequencing (Takara-Clontech) per manufacturer's protocol. cDNA was fragmented using a Covaris E220 sonicator using peak incident power 18, duty factor 20%, cycles per burst 50 for 120 seconds. cDNA was blunt ended, had an A base added to the 3' ends, and then had Illumina sequencing adapters ligated to the ends. Ligated fragments were then amplified for 12-15 cycles using primers incorporating unique dual index tags. Fragments were sequenced on an Illumina NovaSeq-6000 using paired end reads extending 150 bases. Basecalls and demultiplexing were performed with Illumina’s bcl2fastq software and a custom python demultiplexing program with a maximum of one mismatch in the indexing read. RNA-seq reads were then aligned to the Ensembl release 76 primary assembly with STAR version 2.5.1a(*65*). Gene counts were derived from the number of uniquely aligned unambiguous reads by Subread:featureCount version 1.4.6-p5(*66*). Isoform expression of known Ensembl transcripts were estimated with Salmon version 0.8.2(*67*). Sequencing performance was assessed for the total number of aligned reads, total number of uniquely aligned reads, and features detected. The ribosomal fraction, known junction saturation, and read distribution over known gene models were quantified with RSeQC version 2.6.2(*68*).

All gene counts were then imported into the R/Bioconductor package EdgeR(*69*) and TMM normalization size factors were calculated to adjust for samples for differences in library size. Ribosomal genes and genes not expressed in the smallest group size minus one samples greater than one count-per-million were excluded from further analysis. The TMM size factors and the matrix of counts were then imported into the R/Bioconductor package Limma(*70*). Weighted likelihoods based on the observed mean-variance relationship of every gene and sample were then calculated for all samples with the voomWithQualityWeights(*71*). The performance of all genes was assessed with plots of the residual standard deviation of every gene to their average log-count with a robustly fitted trend line of the residuals. Differential expression analysis was then performed to analyze for differences between conditions and the results were filtered for only those genes with Benjamini-Hochberg false-discovery rate adjusted p-values less than or equal to 0.05.

The R EnhancedVolcano package were utilized to generate volcano plots with abs(log2FC) >0 and p-values < 0.05 as cutoff. Gene set-based analyses were performed by entering DEGs [abs(log2FC) >0 and p-values < 0.05] into the ConsensusPathDB (<http://cpdb.molgen.mpg.de/MCPDB> )(*72*) for over-representation analysis in KEGG and Reactome databases with minimal overlap in gene list = 2 and p-value cut-off as 0.01. Heatmaps were generated using Phantasus(*73*).

*Glucose tolerance test (GTT)*

Mice were fasted for 16h before GTT. On the day of GTT, the baseline glucose level of mice was measured by glucometer with a glucose strip with tail blood. Mice were subsequently injected 2mg glucose per gram body weight intraperitoneally (10uL 20% glucose prepared in ddH_2_O, filtered). Blood glucose level was then measured 30min, 60min, and 120min after initial glucose injection. Mice were re-fed after the conclusion of GTT.

*Plasma ALT measurement*

Animal blood was collected from the inferior vena cava into a microtainer with EDTA. Serum was collected after spinning the blood down at 5000rpm for 10min at 4°C and stored at -80°C until assay. ALT quantification was performed using the Teco ALT reagent set (Teco Diagnostics, A524-150) following the manufacturer’s instructions. Briefly, reagents A and B were mixed 5:1 (per sample) to create a working reaction solution and incubated for 10 min at 37°C. Reaction solution (100µL) was then added to 5µL of animal serum (in duplicates) and absorbance at 340nm was read immediately with a TECAN Infinite 200 pro plate reader at 37°C once per minute for 10 min. The average absorbance per minute was determined and multiplied by a factor of 1768 for results in U/L.

*Triglyceride and cholesterol quantification*

Plasma and tissue used for triglyceride or cholesterol quantification originated from animals fasted for 4h before tissue harvest. Tissue was homogenized in cold PBS such that the final homogenate solution contains 100mg tissue/mL. Homogenate solution was diluted 10-times for triglyceride measurement. Following dilution, lipid was solubilized with 1% sodium deoxycholate at 37°C for 5min. Lipid was measured using Infinity Triglyceride Reagent or Cholesterol Reagent (ThermoFisher Cat# TR22421, TR13421) according to the manufacturer’s protocol.

*Lipidomic*

Frozen liver samples (100 to 300mg) were homogenized in water (1:4 w/v) using Omni bead ruptor. Modified Bligh-Dyer method was performed to extract triglyceride (TAG) from 50μL of homogenate after the addition of TAG (17:1/17:1/17:1) as internal standard. Quality control (QC) samples were prepared by pooling the aliquots of the study samples and were used to monitor the instrument stability. The QC was injected six times in the beginning to stabilize the instrument, and was injected between every 5 study samples. Only the lipid species with CV < 15% in QC sample were reported. The relative quantification of lipids was provided, and the data were reported as the peak area ratios of the analytes to the internal standard. Measurement of TAG was performed with a Shimadzu 10A HPLC system and a Shimadzu SIL- 20AC HT auto-sampler coupled to a 4000QTRAP mass spectrometer operated in positive multiple reaction monitoring mode. Data processing was conducted with Analyst 1.6.3.

*Immunofluorescence*

Right lateral lobes were fixed in 10% formalin overnight, washed, then submerged in 30% sucrose (in PBS) and stored at 4°C. For cryo-sectioning, liver lobes were washed in PBS and embedded in OCT compound. Eight µm-thick sections were cut using a cryostat (Leica Biosystems; Wetzlar, Germany) and stored at -80°C until staining. Immunofluorescence staining was carried out as previously published(*26, 64*). Briefly, sections were air-dried for 12min, rehydrated in PBS for 5min, and then blocked in fresh blocking buffer (PBS + Triton X-100 + BSA) for 1h at room temperature. Sections were circled by hydrophobic pen, and primary antibodies prepared in blocking buffer were then deposited onto the section for overnight incubation at 4°C. Next day, sections were washed in PBS 3 times, 5min each, and secondary antibodies were deposited and incubated for 1h at room temperature, in the dark. Sections were washed in PBS 3 times, 5min each. Nuclei were stained with fresh Hoechst dye at 1:25,000 for 5min at room temperature in the dark. Sections were then mounted with prolong gold antifade reagent. Click-iT Plus TUNEL assay kit (AF647) was utilized to detect dying KCs. Confocal images were acquired using an LSM 700 or 900 laser scanning confocal microscope (ZEISS; Jena, Germany) with 10x 0.48 N.A, 20x 0.8 N.A. or 63x 1.4 oil DIC objective at ambient temperature. Samples within an experiment were stained and acquired using the same laser intensity and gain (within the same magnification) on the same day with 4x averaging. Image brightness was uniformly optimized in Fiji (ImageJ, National Institutes of Health, Bethesda, MD, USA).

*qPCR analysis*

For sorted KCs, RNA was purified using the QIAGEN micro RNA kit. For other cell types, Invitrogen mini kits were used. For liver tissue, QIAGEN fibrotic tissue RNA kit was utilized to extract RNA from ~30mg of frozen liver tissue. For cell culture, QIAGEN mini RNA kit was used. RNA concentration was determined by nanodrop spectrophotometer. Reverse transcription was performed using the High-Capacity cDNA Reverse Transcription Kit (Applied Biosystems). qPCR was performed using Power SYBR green reagent (Applied Biosystem) with a Quant Studio 3 platform. Relative gene expression was calculated using delta-delta CT methods and normalized to *36b4* expression. qPCR primers used is listed in Supplemental Table 1.

*Histology*

Right lobes were fixed in 10% formalin at 4°C overnight and then transferred into 70% ethanol for long-term storage at 4°C. Paraffin-embedding, sectioning, H&E, and picrosirius red staining (PSR) were performed by the Advanced Imaging and Tissue Analysis Core of the Digestive Disease Research Core Center at Washington University in St. Louis. Steatosis scores were quantified by a blinded liver pathologist using previously described criteria(*74*). Macrovesicular and microvesicular steatosis were scored from 0 to 3 and quantified as percentages (<5%, grade 0; 5-33%, grade 1; 34-66%, grade 2; >66% grade 3). Picrosirius red area was quantified blinded with ImageJ using manual thresholding on 7 images at high-power magnification per sample (randomly chosen area avoiding major vessels). Images were taken on an AxioImager M.2.

*Plasma ELISA*

Plasma GDF15 was measured using the mouse GDF15 DuoSet ELISA kit (R&D systems, Cat# DY6385) following manufacturer’s instruction.

*Seahorse cell mitostress test*

BMDMs were seeded at 5 x 10^5^ cells/mL onto Agilent Seahorse XF96 cell culture microplate for 3h to allow adhesion. The Mitostress test was performed according to the manufacturer’s protocol for Seahorse XF Cell Mito Stress Test. Briefly, 1.3µM Oligomycin (Cayman chemical; Cat #11341), 1µM FCCP (Sigma; Cat #C2920), and 1µM rotenone (Sigma; Cat #8875) together with 10µM antimycin-A (Sigma A-8674) were added to BMDMs every 18 minutes. Oxygen consumption rate (OCR) and extracellular acidification rate (ECAR) were recorded.

*In vitro lipid accumulation assay*

For assay with free fatty acid (FA), BMDMs were plated on non-tissue culture-treated plates or on #1.5 coverslips overnight, then incubated with 250µM oleic acid conjugated with BSA (2:1 molar ratio) and 1µM of BODIPY-C_16_ in 5% CMG-containing DMEM. After incubation for indicated time points, cells were washed with PBS, trypsinized, and resuspended with 1:3000 DAPI solution for flow cytometry. For imaging, cells were fixed with 4% PFA for 15 min at room temperature in the dark, followed by thorough washing with PBS. Nuclei were stained with 1:25000 Hoechst dye for 5min at room temperature in the dark. Coverslips were then mounted to slides with prolong gold antifade reagent and stored in dark.

*Cell death and ROS measurement in cell culture*

BMDMs were plated on suspension plates and subsequently treated with various cell death inducers in complete DMEM without additional CMG-conditioned media. RSL3 (5µM, Selleckchem, Cat#S8155) or ML162 (5µM, Cayman Chemical, Cat#20455) and ferrostatin-1 (5µM, Cayman Chemical, Cat#17729) were used to induce or inhibit ferroptosis, respectively. zVAD(OMe)-FMK (20µM, Santa Cruz, Cat#SC-311561) + LPS (100ng/mL, in PBS) and necrostatin-1 (20µM, EMD-Calbiochem, Cat#4311-88-0) were used to induce or inhibit necroptosis, respectively. DMSO was used as vehicle control. For H_2_O_2_ experiments, cells were treated with 2.5mM H_2_O_2_ for 2h; water was used as vehicle control. To inhibit ACLY and SCD1, cells were pretreated with BMS303141 (ACLYi) (40µM) or CAY10566 (SCD1i) (10nM) respectively for 16h, and then co-treated with RSL3 (5µM) for 3h followed by lipid peroxidation measurement. For lipid peroxidation measurement, BODIPY 581/591 C_11_ (Invitrogen, Cat# D3861) were incubated with cells at a final concentration of 5µM for 30min at 37°C; for cellular oxidative stress measurement, CellRox Deep Red was used at 1µM (in serum-free DMEM) and incubated with cells for 30min at 37°C. Cell death was measured by adding propidium iodide solution (1:500 in FACS buffer) to cells after indicated time points. Data were acquired with a BD Canto II cytometer.

*NADP^+^ and NADPH measurement*

NADP+ and NADPH in BMDMs and KCs were measured by using Promega NADP/NADPH-Glo™ Assays (G9081), following manufacturer’s instruction. Briefly, cells were suspended in 80µL PBS per well and lysed with equal volume of base solution containing 1% DTAB. Lysate from each sample (50µL) was transferred into empty PCR tubes containing either nothing (base-treated samples) or 25µL 0.4N HCl (acid-treated samples) and heated at 60C on a PCR block. After heating, samples were equilibrated to room temperature for 10 min. To the acid-treated samples, 25µL 0.5M Trizma base was added; to base-treated samples, 50µL 0.2N HCl/0.25M Trizma solution was added. To a 96-well white-wall assay plate, 50µL of acid- or base-treated samples were transferred, and equal volume of NADP^+^/NADPH-glo detection reagent was added and incubated for 30min at room temperature. Luminescence was detected and recorded by a TECAN Infinite 200 pro plate reader. For BMDM assays, 7x10^5^ cells/mL cells in quadruplicates plated on 96-well plate was used. For KC assays, primary murine KCs were isolated as described in “Cell culture” section, with the additional step of normalizing the number of CD45^+^ cells to 1x10^6^ cells/mL for plating onto 96-well plates in triplicates. Two hours after plating, non-adherent cells were removed and adherent cells (KCs) were lysed following assay instruction.

*Metabolite measurement from cells*

BMDMs were washed twice with PBS, twice with LC-MS grade water, and then quenched and collected with ice-cold LC-MS grade methanol into Eppendorf tubes. Cells were dried in a SpeedVac for 2h, and then reconstituted in 1mL of col methanol:acaetonitrile:water in 2:2:1 ratio. Cell suspension was then vortexed, frozen in liquid nitrogen, and bath sonicated for 10min at 25°C, for a total of 3 times. Samples were then stored at -20°C for 1h, and centrifuged at 14,000 rpm at 4°C for 10min. Supernatants were transferred to new Eppendorf tubes and dried b SpeedVac for 2h, while pellets were resuspended in 200µL of 50mM sodium hydroxide for protein quantification using BCA assay (ThermoFisher). To the dried residues, 1µL of water:acetonitrile in 1:2 ratio was added for every 2.5µg of protein. The samples were then bath sonicated for 5min at 25°C and vortexed, for a total of 2 times, and stored at 4°C for 1h. Then samples were centrifuged at 14000 rpm at 4°C for 10min. Pellets were discarded and supernatants were transferred into LC vials and stored at -80°C until mass spectrometry analyses.

*Statistics*

All statistical analyses were performed using Prism 10.3 software. Two-tailed, unpaired t-tests were used when 2 independent groups were being compared. One-way ANOVA followed by Šídák’s multiple comparison tests were utilized for when more than 2 independent groups were being compared with each other or with a control group. Two-way ANOVA followed by multiple t-tests were performed whenever appropriate. Specific statistical tests, biological replicates (n) were described in figure legends. P-value ≤ 0.05 was considered statistically significant for all analyses.

*Antibodies used for flow cytometry*

| Target | Conjugate | Clone | Suppliers | Catalog # | Dilution |
| --- | --- | --- | --- | --- | --- |
| CD45 | BUV395 | 30-F11 | BD Biosciences | 564279 | 1:100 |
| CD45 | PerCP-Cy5.5 | 30-F11 | BioLegend | 103132 | 1:100 |
| CD11b | APC-Cy7 | M1/70 | BioLegend | 101226 | 1:100 |
| CLEC2 | APC | 17D9/CLEC-2 | Biolegend | 146105 | 1:100 |
| CLEC2 | PE | 17D9/CLEC-2 | Biolegend | 146103 | 1:100 |
| F4/80 | APC | BM8 | BioLegend | 123122 | 1:100 |
| F4/80 | FITC | BM8 | BioLegend | 123107 | 1:100 |
| F4/80 | BV605 | BM8 | BioLegend | 123133 | 1:100 |
| TIM4 | BV421 | 21H12 | BD Biosciences | 742773 | 1:500 |
| TIM4 | PE-Cy7 | RMT4-54 | BioLegend | 130010 | 1:100 |
| VSIG4 | PE-Cy7 | NLA14 | Invitrogen | 25-5752-80 | 1:200 |
| VSIG4 | FITC | NLA14 | Invitrogen | 53-5752-82 | 1:200 |
| MHCII | BV605 | M5/114.15.2 | BioLegend | 107639 | 1:300 |
| Ly6C | BV711 | HK1.4 | BioLegend | 128037 | 1:100 |
| Ki67 | BV605 | 16A8 | BioLegend | 652413 | 1:100 |

*Antibodies used for immunofluorescence*

| **Species** | **Target Species** | **Antigen** | **Clone** | **Suppliers** | **Cat. #** | **Dilution** |
| --- | --- | --- | --- | --- | --- | --- |
| Rat | Mouse | F4/80-Biotin | BM8 | eBioscience | 13-4801-85 | 1:200 |
| Rat | Mouse | CD68 | FA-11 | Invitrogen | 14-0681-82 | 1:200 |
| Rat | Mouse | TIM4 | RMT4-54 | BioLegend | 130002 | 1:100 |
| Goat | Mouse | CLEC4F/CLECSF13 | Polyclonal | R&D Systems | AF2784 | 1:100 |

| **Host Species** | **Target Species** | **Target** | **Conjugate** | **Suppliers** | **Cat. #** | **Dilution** |
| --- | --- | --- | --- | --- | --- | --- |
| Donkey | Rat | IgG(H+L)_ | AF 488 | Invitrogen | A21208 | 1:500 |
| Donkey | Goat | IgG (H+L) | AF 647 | Jackson ImmunoResearch |  | 1:500 |
| Donkey | Rabbit | IgG (H+L) | AF594 | Jackson ImmunoResearch | 711-585-152 | 1:200 |
| Donkey | Rabbit | IgG (H+L) | AF647 | Jackson ImmunoResearch | 711-605-152 | 1:500 |

**Fig. S1. Gating strategy for hepatic myeloid cells, Clec4f^Cre^ (KC^Cre^) validation, and baseline functional assessment of TFEB-KCs.** (**A**) Gating strategy for identifying liver myeloid cells and monocytes in HFHS and STD-fed mice. (**B**) TdTomato reporter signal and representative histogram in various liver myeloid cells in KC^Cre-TdT^, KC^Tfeb-TdT^, and no Cre control (Cre^neg^TdT^pos^) mice fed 16 weeks of HFHS diet (n = 8-10/group). (**C-D**) KCs were isolated from female KC^Cre^ and KC^Tfeb^ mice and incubated with various substrates to measure macrophage function and lysosomal activity (n = 5/group). (C) Schematic of experiments. (D) MFI of TMR dextran, lysotracker green, pHrodo, and DQ-OVA in WT and TFEB-KCs. Data represents individual biological replicates and are presented as means ±SEM. P-values were calculated using (B) two-way ANOVA followed by multiple t-tests, and (D) unpaired two-tailed Student’s t-tests. NS = not significant, *p < 0.05, **p < 0.01, ***p< 0.001, ****p<0.0001.

**Fig. S2. Systemic obesity of male and female KC^Cre^ and KC^Tfeb^ mice fed MASLD/MASH-inducing diets.** (**A**) Kinetics of weight gain for male and female KC^Cre^ and KC^Tfeb^ mice during HFHS diet feeding (n = 6-8 for females, and 26-29 for males). (**B**) Glucose tolerance test (GTT) and quantified area under the curve (AUC) for male KC^Cre^ and KC^Tfeb^ mice fed 16 weeks HFHS diet (n = 7-8 per group). # represents comparisons between STD- and HFHS-fed KC^Tfeb^ mice and * represents comparisons between STD- and HFHS-fed KC^Cre^ mice. (**C**) Flow cytometric quantification of adaptive immune cells (CD3^+^CD4^+^T cells, CD3^+^CD8^+^T cells, and CD3^-^B220^+^ B cells), in KC^Cre^ and KC^Tfeb^ mice fed 16-week HFHS diet. (**D-I**) KC^Cre^ and KC^Tfeb^ male mice were fed 8 weeks of HFHS diet (n = 7-9/group). (D) Final body and organ weights of mice. (E) Flow cytometric quantification of liver macrophages per gram of tissue. (F) KC number per gram of tissue and as a percentage of liver macrophages. (G) MdM number, (H) MoKC, LAMs, and (I) Ly6C^hi^ monocytes per gram of tissue. (**J-O**) Female KC^Cre^ and KC^Tfeb^ mice were fed HFHS diet for 16 weeks (n = 6-8/group). (J) Final body and organ weights of mice. (K) Flow cytometric quantification of liver macrophages per gram of tissue. (L) KC number per gram of tissue and as a percentage of liver macrophages. (M) MdM number, (N) MoKC, LAMs, and (O) Ly6C^hi^ monocytes per gram of tissue. (**P**) Gating strategy for myeloid cells found in livers of mice fed CDAA diet utilizing CLEC2. Data represents (A, B) means ±SEM or (B-O) individual biological replicates presented as means ±SEM. P-values were calculated using (A) unpaired two-tailed Student’s t-tests between each time point, (B) two-way ANOVA followed by multiple t-tests, and (C-O) unpaired two-tailed Student’s t-tests. NS = not significant, *p < 0.05, **p < 0.01, ***p< 0.001, ****p<0.0001.

**Fig. S3. Characterization of steatosis and fibrosis in KC^Cre^ and KC^Tfeb^ mice.** (**A**) Quantification of hepatic cholesterol (n = 18-22/group) and serum triglyceride (n = 2-16/group) in KC^Cre^ and KC^Tfeb^ male mice fed 16 weeks of STD or HFHS diet (n = 18-22/group). (**B-D**) KC^Cre^ and KC^Tfeb^ male mice fed 16 weeks of STD or HFHS diet and livers were used for lipidomic. (B) Body parameters and KC preservation of mice whose livers were subjected to targeted lipidomic (n = 3-6/group). (C) TAG species with significant changes between KC^Cre^ and KC^Tfeb^ mice measured by lipidomic. (D) qPCR gene expression analyses of pathogenic collagens in whole liver tissues (n = 5-9/group). (**E**) Representative PSR staining of livers from CDAA diet-fed KC^Cre^ and KC^Tfeb^ mice and quantification of PSR^+^ % by averaging signal from 7 images per mouse (n = 5/group). Scalebar = 100µm. Data represents individual biological replicates presented as means ±SEM. P-values were calculated using (A, F) unpaired two-tailed Student’s t-tests and (B-E) two-way ANOVA followed by multiple t-tests. NS = not significant, *p < 0.05, **p < 0.01, ***p< 0.001, ****p<0.0001.

**Fig. S4. Transcriptomic pathways altered by TFEB and HFHS diet in KCs.** (**A-B**) DEGs in TFEB-KCs under homeostatic condition. (A) Volcano plot showing DEGs and (B) upregulated KEGG pathways in TFEB-KCs. (**C**) Venn diagram of upregulated DEGs between WT and TFEB-KCs induced by HFHS. (**D**-**E**) DEGs in WT-KCs after HFHS diet feeding. (D) Volcano plot showing DEGs and (E) differential KEGG pathways induced by HFHS in WT-KCs. (**F-G**) DEGs in TFEB-KCs after HFHS diet feeding. (F) Volcano plot showing DEGs and (G) differential KEGG pathways induced by HFHS in TFEB-KCs. (**H**) Heatmap and p-values of KC2 signature identified by Blériot et al., 2021(*25*).

**Fig. S5. Lipid accumulation and metabolic requirement in TFEB-KCs.** (**A-D**) C57BL/6 male mice were fed STD or HFHS diet for 16 weeks. (A) MFI of BODIPY-C_16_ signal in STD or HFHS KCs by flow cytometry after 1 minute incubation with FA (n = 5/group). (B) Representative electron micrographs of FACS-purified KCs, scalebar = 2µm. (C) Total number of LDs, (D) distribution of cells with 0-2, 3-5, or 5+ LDs, (E) total LD area, and relative area of LD with respect to total cytosol area across 20 cells per condition. (**F**) Seahorse mitostress test on WT or TFEB-BMDMs. Oligo: oligomycin; Rot/Ant: rotenone/antimycin; OCR: oxygen consumption rate. (**G**) qPCR gene expression analyses of *Cpt2* and *Lipa (*encodes for LAL) in KCs isolated from respective mouse lines (n = 2-4/group). (**H-K**) Male mice fed 16-week HFHS diet. Body parameters, organ weights, and liver triglycerides of (H) KC^Cre^CPT2^fl/fl^ and KC^Tfeb^CPT2^fl/fl^ mice (n = 6-16/group), and (I) KC^Cre^LAL^fl/fl^ and KC^Tfeb^LAL^fl/fl^ (n = 5-7/group) (J-K) H&E images of livers from mice in fig. S5H and S5I . Scalebar = 500µm. (**L**) Serum GDF15 measured by ELISA from Mac^Cre^ and Mac^Tfeb^ mice fed 16-week STD or high-fat diet (HFD) (n = 3-6/group). (**M**) qPCR gene expression analyses of *Gdf15* in KCs and BMDMs isolated from respective mouse lines (n = 2-3/group). (**N-P**) Male KC^Cre^GDF15^fl/fl^ and KC^Tfeb^GDF15^fl/fl^ were fed 16-week of HFHS diet (n = 7-9/group). (N) Body and organ weights of mice. (O) H&E images of livers. (P) Liver TAG measurement (n = 6-7/group). Data represents (A, G-I, L-N, P) individual biological replicates or (F) technical triplicates presented as means ±SEM. P-values were calculated using (A, F-I, M-N, P) unpaired two-tailed Student’s t-tests and (L) two-way ANOVA followed by multiple t-tests. NS = not significant, *p < 0.05, **p < 0.01, ***p< 0.001, ****p<0.0001.

**Fig. S6. Loss of TFEB in macrophages, and functional characterization of KCs during MASLD.** (**A-D**) Control (TFEB^fl/fl^) and LysM-Cre-driven TFEB-deficient mice (Mac^Cre^TFEB^fl/fl^) at homeostatic condition and after 8 weeks of CDAA diet. Circles represent males and triangles represent females. (A) qPCR gene expression analyses of *Tfeb* and (B) *Tfe3* in BMDMs. (C) Flow cytometric quantification of myeloid subsets, including KCs, (D) MdMs, and Ly6C^hi^ monocytes (n = 4/group). (**E**-**F**) KCs were isolated from C56BL/6 male mice fed 16 weeks of STD or HFHS diet and incubated with various substrates to measure lysosomal activity (n = 5/group). (E) Flow cytometry quantification of KCs per gram of tissue. (F) MFI of lysotracker Green, pHrodo, and DQ-OVA in KCs. (**G**) MFI of Lysotracker Green and DQ-OVA in KCs isolated from KC^Cre^ and KC^Tfeb^ male mice fed 16 weeks of STD or HFHS diet (n = 2-3/group). (**H-L**) C57BL/6 male mice were fed STD or HFHS for 16 weeks and livers were *in situ* injected with fluorescent beads (n = 4-5/group). (H) Schematic of experiment. (I) Representative flow histogram of bead signal in KCs. (J) Percentage of KCs with single or multiple bead positive signals. (K) Representative immunofluorescence images of bead capturing in KCs. Red: beads; green: TIM4; blue: DAPI. Scale bar = 30µm. (L) Flow cytometric quantification of MdMs per gram of tissue. (M) Percentage of myeloid cells and monocytes with bead positive signal. (**N**) Quantification of KCs per gram of tissue and as percentage in KC^Cre^ and KC^Tfeb^ mice used for in situ fluorescent bead assay. Refers to main Fig. 6J-K. Data represents individual biological replicates and are presented as means ±SEM. P-values were calculated using (A-F,J, L, N) unpaired two-tailed Student’s t-tests, (G) two-way ANOVA followed by multiple t-tests, and (M) one-way ANOVA followed by multiple t-tests. NS = not significant, *p < 0.05, **p< 0.01, ***p< 0.001, ****p<0.0001.

**Fig. S7. Loss of MdMs in KC^Tfeb^ mice with additional knockout, macrophage cell death and gene expression analyses.** (**A**) Flow cytometric quantification of myeloid and monocyte subsets in the livers of male and female KC^Cre^CPT2^fl/fl^ and KC^Tfeb^CPT2^fl/fl^ mice after 16 weeks of HFHS diet (n = 15-18/group) or 8 weeks of CDAA diet feeding (n = 8-10/group). (**B**) Flow cytometric quantification of myeloid and monocyte subsets in the livers of male and female KC^Cre^GDF15^fl/fl^ and KC^Tfeb^ GDF15^fl/fl^ mice after 16 weeks of HFHS diet (n = 11-12/group) or 10 weeks of CDAA diet feeding (n = 4/group). (**C**) Flow cytometric quantification of myeloid and monocyte subsets in the livers of male and female KC^Cre^LAL^fl/fl^ and KC^Tfeb^ LAL^fl/fl^ mice after 16 weeks of HFHS diet (n = 6-7/group) or 10 weeks of CDAA diet feeding (n = 11/group). (**D**) Ferroptotic death was measured by propidium iodide^+^ (PI^+^) signal in WT- or TFEB-BMDMs treated with 5µM ML162 ± 5µM Fer1 for 2h. (**E**) Flow cytometric percentage of PI^+^ and oxidized BODIPY C11^+^ signal in cells treated with 5µM RSL3 for 1h, 3h, 6h, 12h, and 24h. Solid symbols represent oxidized BODIPY-C_11_ signal and open symbols represent PI staining. (**F**) Secreted IL-1β in WT- or TFEB-BMDMs stimulated with LPS and ATP. N.D. = not detectable. (**G-H**) Gene expression analyses on (G) antioxidant enzymes and (H) key enzymes in the PPP in WT- or TFEB-BMDMs. (**I**) Ribose-5-phosphate level in WT- and TFEB-BMDMs measured by mass spectrometry. (**J**) Lipid peroxidation was measured by oxidized BODIPY-C_11_ signal in WT-BMDMs pre-treated with ACLY inhibitor (BMS 303141) or SCD1 inhibitor (CAY10566) followed by co-treatment with RSL3 for 3h. (**K**) qPCR analysis of PPP-related genes in primary KCs (n = 3-4/genotype). Data represents (A-C, J) individual biological replicates, (D-F, I) technical triplicates and (G-H) 2 biological replicates with technical triplicates presented as means ±SEM. P-values were calculated using (A-C, F-I, K) unpaired two-tailed Student’s t-tests, (D, J) two-way ANOVA followed by multiple t-tests. NS = not significant, *p < 0.05, **p < 0.01, ***p< 0.001, ****p<0.0001.
