## Supplementary figures and images for "Induction of TFEB promotes Kupffer cell survival and reduces lipid accumulation and inflammation in MASLD"

### Supp. Fig 1

##### Supplemental Figure 1. Related to Main Figure 1.

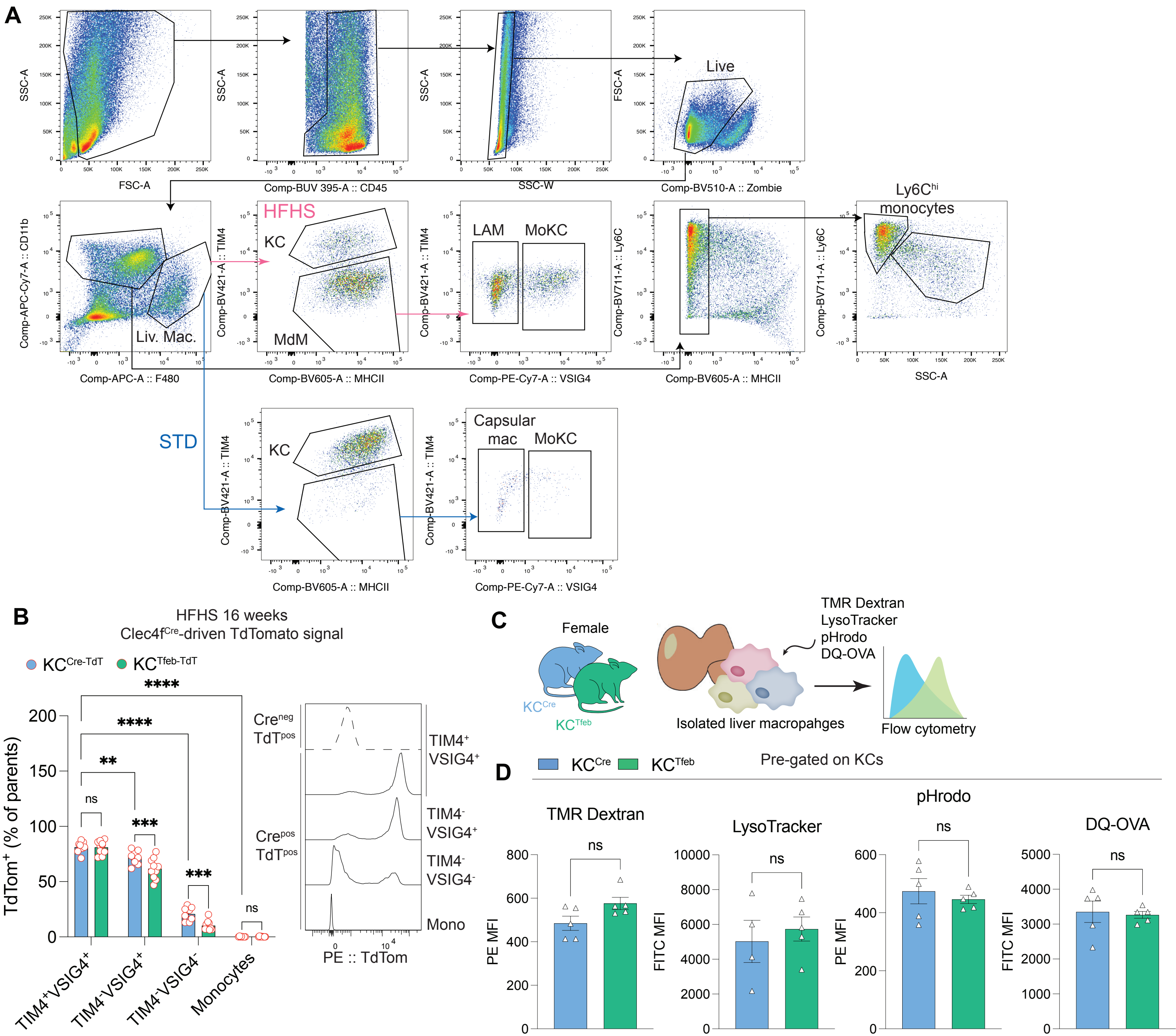

### Supp. Fig 2

#### Supplemental Figure 2. Related to Main Figure 2.

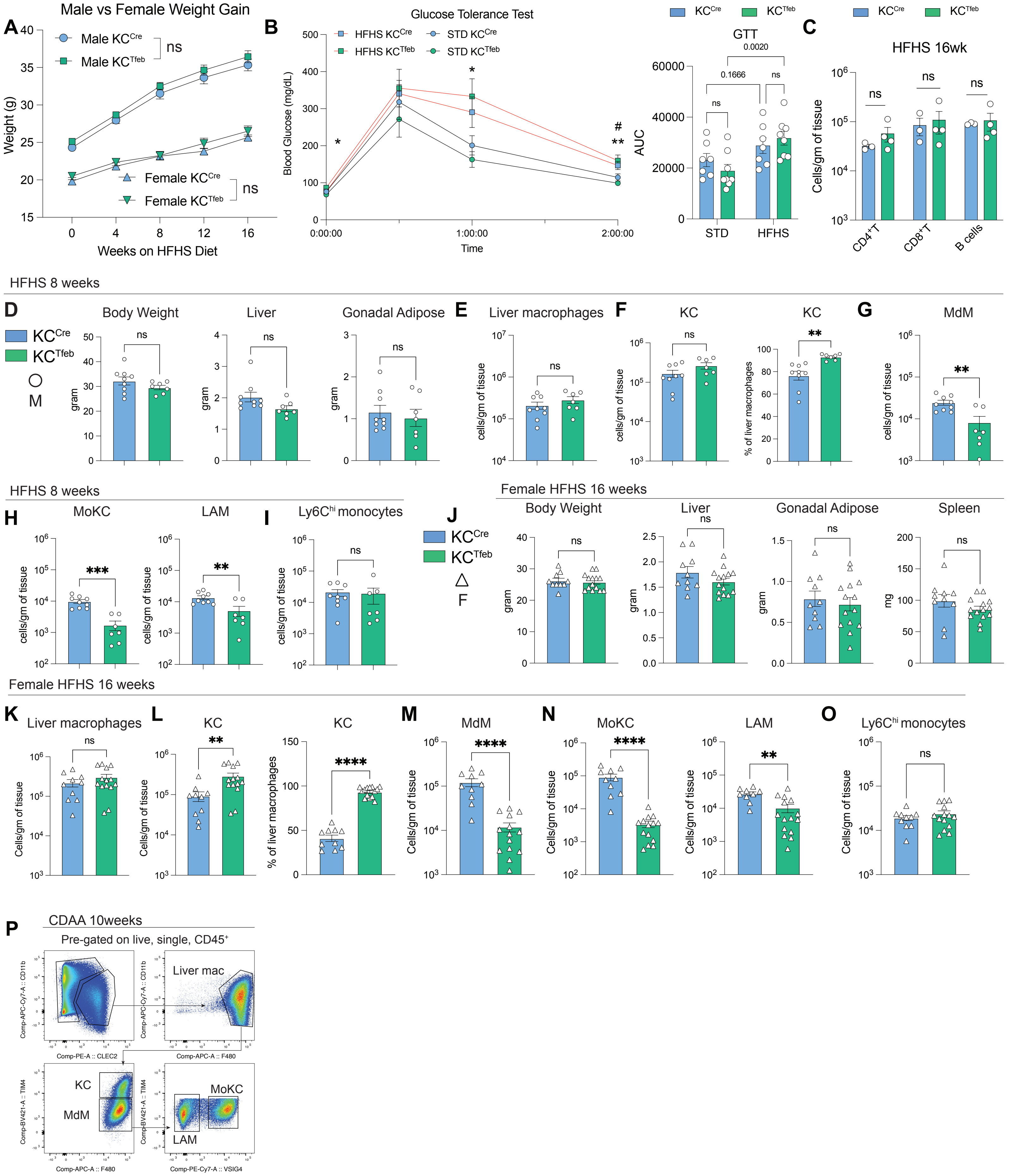

### Supp. Fig 3

#### Supplemental Figure 3. Related to main figure 3.

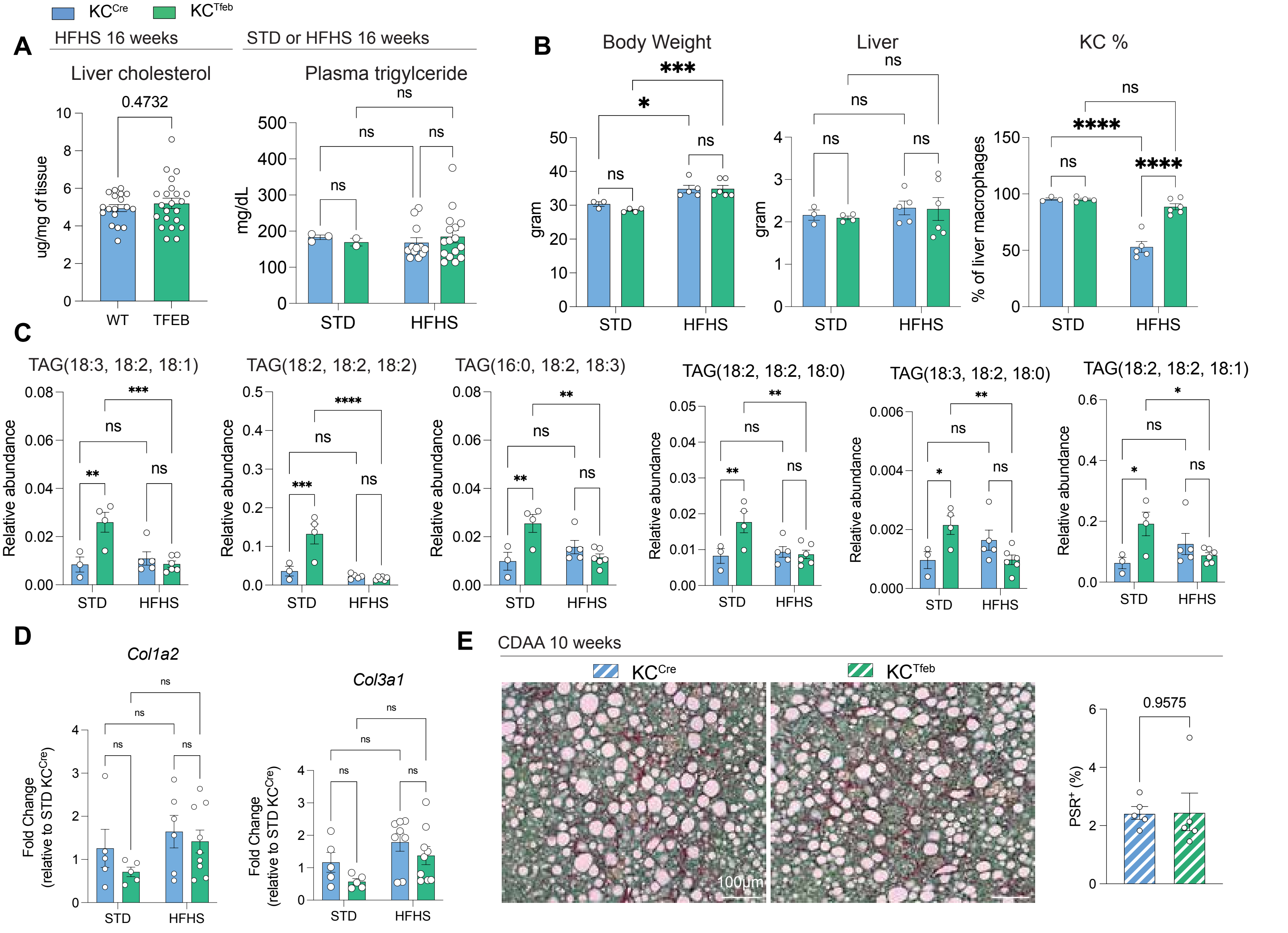

### Supp. Fig 5

##### Supplemental Figure 5. Related to Main Figure 5.

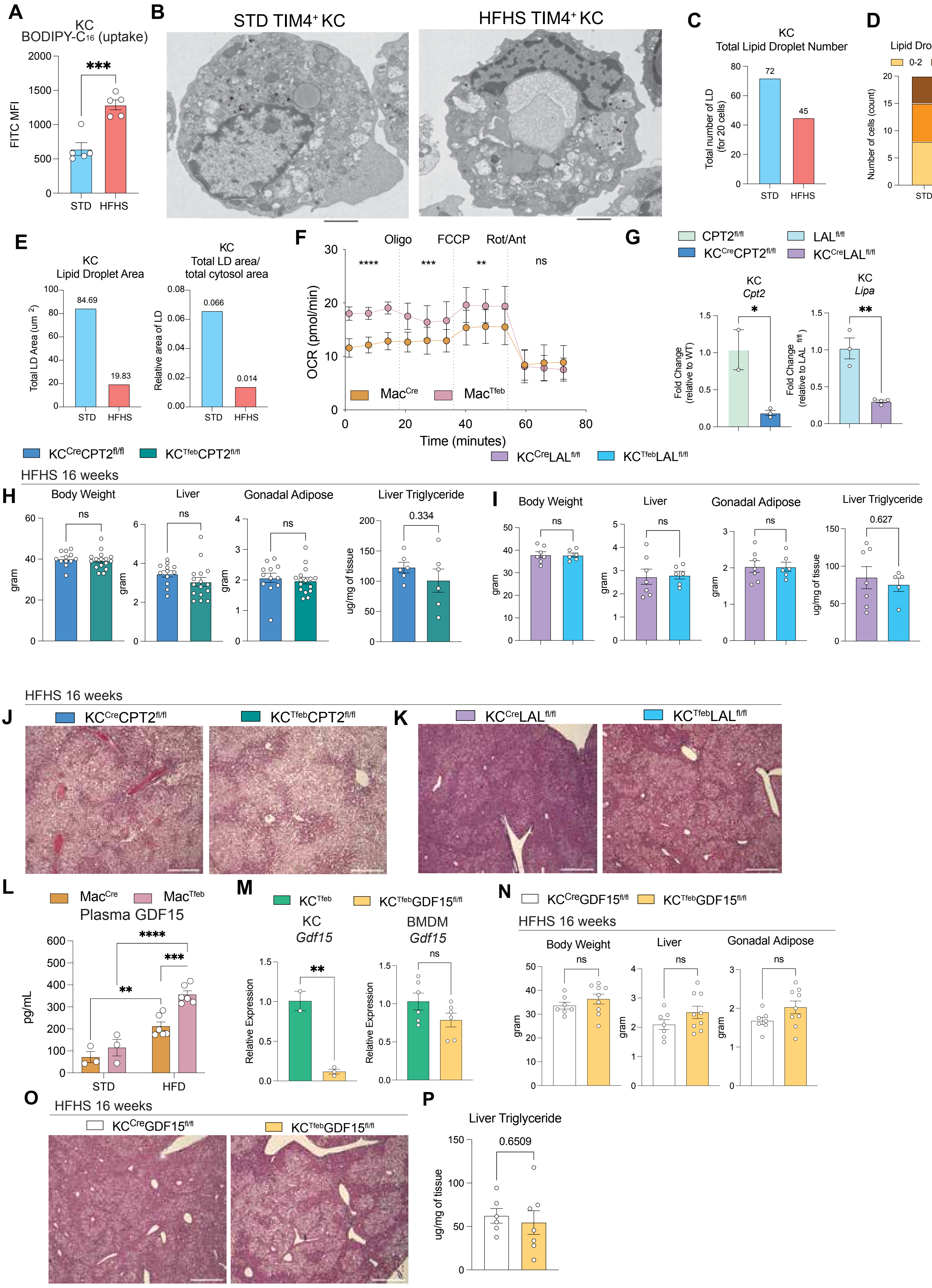

### Supp. Fig 6

#### Supplemental Figure 6. Related to Main Figure 6.

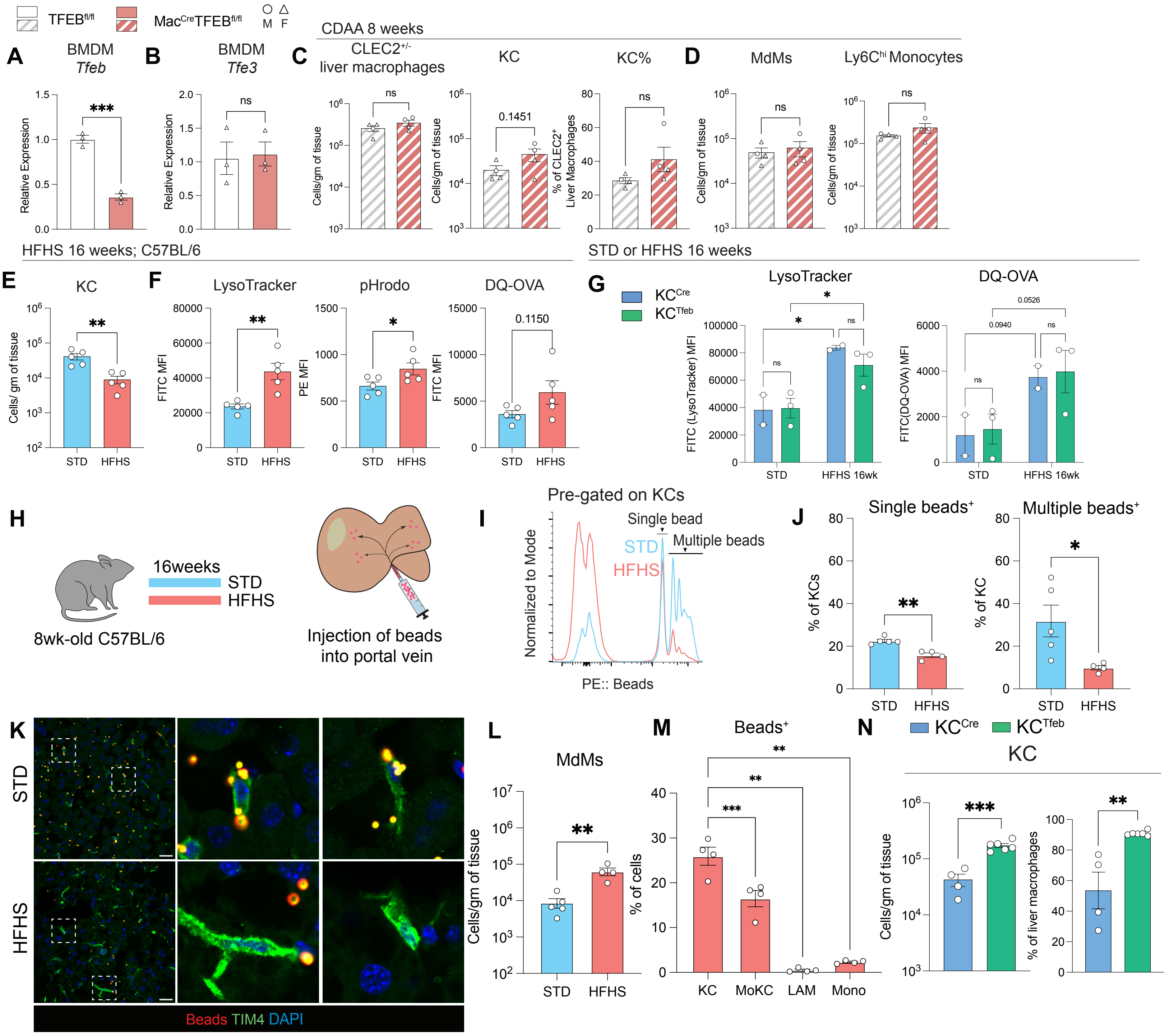

### Supp. Fig 7

#### Supplemental Figure 7. Related to Main Figure 7.

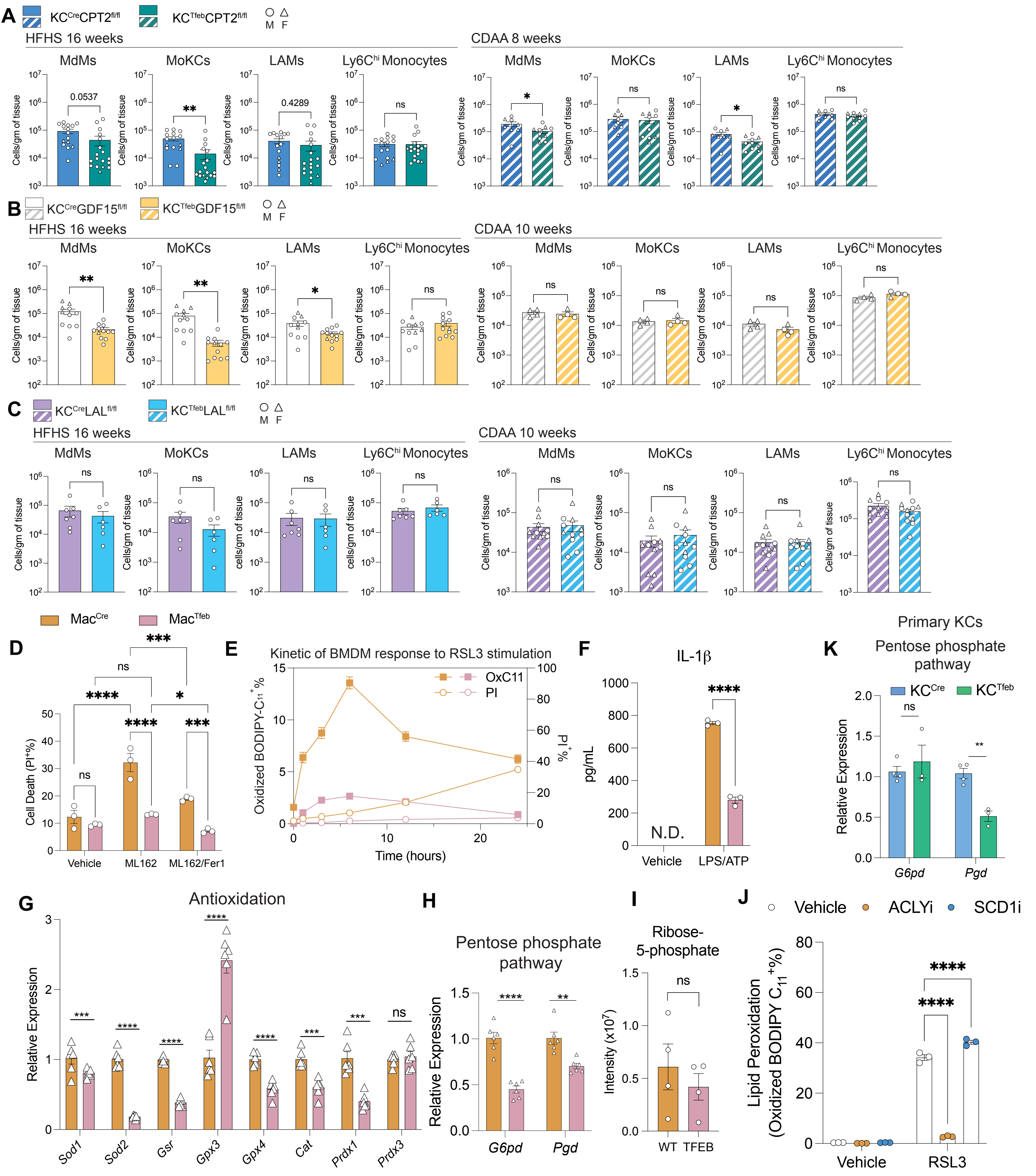
