## Supplementary material for "Induction of TFEB promotes Kupffer cell survival and reduces lipid accumulation and inflammation in MASLD": Supp. Fig 4

### Supplemental Figure 4. Related to main figure 4.

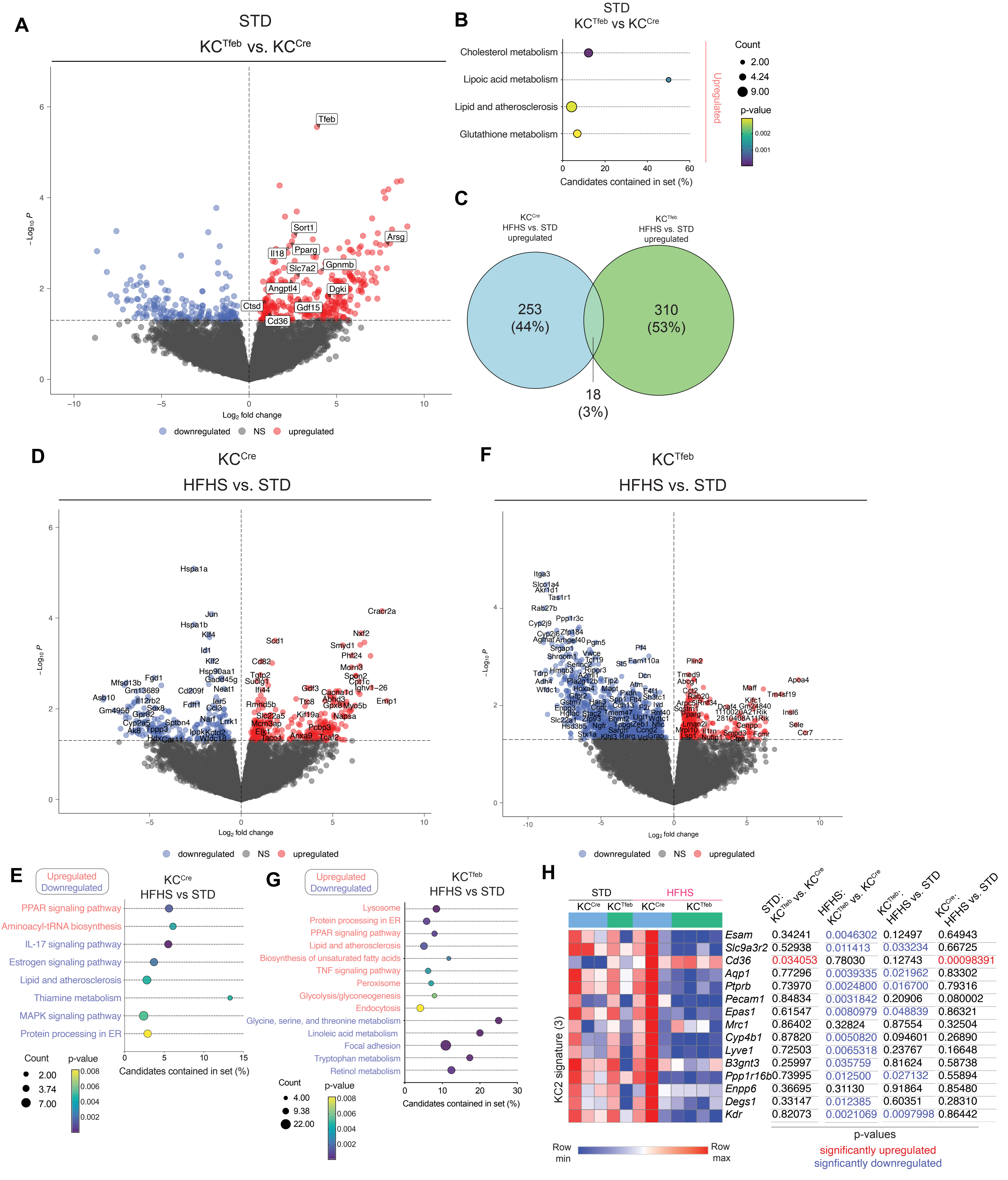


| | STD: KCTfeb vs. KCCre | HFHS: KCTfeb vs. KCCre | KCTfeb: HFHS vs. STD | KCCre: HFHS vs. STD |
|---|---|---|---|---|
| *Esam* | 0.34241 | 0.0046302 | 0.12497 | 0.64943 |
| *Slc9a3r2* | 0.52938 | 0.011413 | 0.033234 | 0.66725 |
| *Cd36* | 0.034053 | 0.78030 | 0.12743 | 0.00098391 |
| *Aqp1* | 0.77296 | 0.0039335 | 0.021962 | 0.83302 |
| *Ptprb* | 0.73970 | 0.0024800 | 0.016700 | 0.79316 |
| *Pecam1* | 0.84834 | 0.0031842 | 0.20906 | 0.080002 |
| *Epas1* | 0.61547 | 0.0080979 | 0.048839 | 0.86321 |
| *Mrc1* | 0.86402 | 0.32824 | 0.87554 | 0.32504 |
| *Cyp4b1* | 0.87820 | 0.0050820 | 0.094601 | 0.26890 |
| *Lyve1* | 0.72503 | 0.0065318 | 0.23767 | 0.16648 |
| *B3gnt3* | 0.25997 | 0.035759 | 0.81624 | 0.58738 |
| *Ppp1r16b* | 0.73995 | 0.012500 | 0.027132 | 0.55894 |
| *Enpp6* | 0.36695 | 0.31130 | 0.91864 | 0.85480 |
| *Degs1* | 0.33147 | 0.012385 | 0.60351 | 0.28310 |
| *Kdr* | 0.82073 | 0.0021069 | 0.0097998 | 0.86442 |
